# Redox oscillation frequency reshapes antibiotic resistance-gene expression through metabolic and ecological selection in a freshwater community

**DOI:** 10.64898/2026.09.29.755287

**Authors:** Bharat Manna, Ishi Keenum, Qian Liu, Naresh Singhal

**Affiliations:** School of Bio Sciences and Technology, Vellore Institute of Technology, Vellore, Tamil Nadu 632014, India; Civil, Environmental, and Geospatial Engineering, Michigan Technological University, Michigan 49931, USA; State Key Laboratory of Satellite Ocean Environment Dynamics, Second Institute of Oceanography, Ministry of Natural Resources, Hangzhou 310012, China; Department of Civil and Environmental Engineering, University of Auckland, Auckland 1142, New Zealand; Water Research Centre, University of Auckland, Auckland 1142, New Zealand

**Keywords:** oxic–anoxic cycling, central metabolism, oxidative stress, metatranscriptomics, strain-resolved metagenomics, efflux pumps, *Pseudomonas*

## Abstract

Aquatic environments can maintain and express antibiotic resistance genes even without antibiotic pressure, but the conditions that govern their activity remain poorly understood. Redox oscillations are pervasive in natural and engineered waters and strongly alter microbial metabolism, yet whether their frequency shapes resistance expression is unknown. We reanalyzed paired metagenomes and metatranscriptomes from a published chemostat experiment in which a sulfidic stream community was cycled between oxic and anoxic conditions at high, medium and low frequency (HF, MF, LF), with strain-resolved population genomics of the dominant *Pseudomonas* lineage. Resistance genes were broadly distributed across treatments, whereas transcription concentrated under fast cycling, reaching a median 168 TPM under HF against 38 under LF although no antibiotic was present, so genetic potential and expression diverged. Fast cycling induced ROS-generating flavoproteins, antioxidant defenses and transcription-coupled DNA repair, and *Pseudomonas* supplied 99% of resistance-gene transcripts under HF, chiefly through its intrinsic efflux systems. Within the dominant *Pseudomonas* genome, however, the four resistance genes carried no sequence variants under HF, where coverage was deepest, and no fixed difference from the reference in any sample. Genome-wide selection remained purifying throughout (pooled pN/pS 0.72 under HF against 0.12 under LF, both below 1), and the relaxation trended toward redox and central metabolism. Redox frequency therefore raises resistance expression by selecting a metabolically versatile, high-efflux lineage, while the resistance sequences themselves stay conserved. Resistance activity in aquatic systems responds to an abiotic driver, which argues for weighing expression alongside gene presence in environmental surveillance.

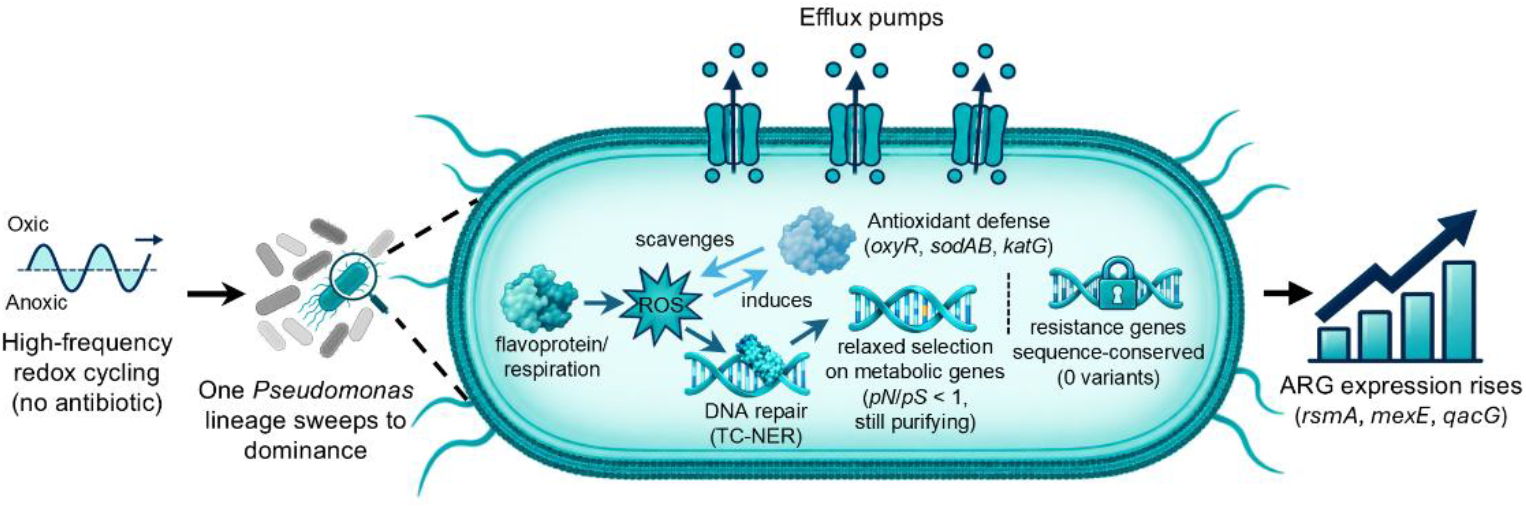

**Highlights:**

- Fast redox cycling raised ARG expression without any antibiotic added.
- High-frequency cycling induced ROS sources, antioxidant defense and DNA repair.
- Fast cycling swept one *Pseudomonas* lineage to community dominance.
- ARGs in the dominant *Pseudomonas* genome carried no variants under HF.
- Gene-based AMR surveillance would miss the rise in resistance expression.

## 1. Introduction

Antimicrobial resistance (AMR) is among the most serious threats to modern medicine, with projections of 10 million annual deaths by 2050 if current trends continue (G B D 2021 Antimicrobial Resistance Collaborators, 2024). Resistance continues to erode the effectiveness of established antibiotics and the care that depends on them (Edwards et al., 2021). Global efforts have focused mainly on limiting antibiotic use through stewardship, yet resistance continues to emerge (Amábile-Cuevas, 2025). A central question is why resistance keeps arising and spreading even when direct antibiotic selection is minimized (Andersson and Hughes, 2012). This persistence indicates that the bacterial evolution underlying AMR proceeds through mechanisms more varied than antibiotic selection alone (Bakkeren et al., 2020; Koolman et al., 2025), and that management focused on antibiotic exposure addresses only part of the process (Knöppel et al., 2017).

Recent work has broadened this picture, showing that resistance can evolve through mechanisms extending well beyond direct antibiotic selection (Souque et al., 2024). Lopatkin *et al*. found that clinically relevant resistance can arise through mutations in core metabolic genes, including 2-oxoglutarate dehydrogenase (*sucA*), isocitrate dehydrogenase (*icd*), and glutamate synthase (*gltB, gltD*), rather than through canonical resistance targets. These mutations lower basal respiratory activity and limit antibiotic-induced metabolic toxicity, and they are enriched across more than 3,500 pathogenic *E. coli* isolates (Lopatkin et al., 2021). Knöppel *et al*. showed that *E. coli* and *Salmonella enterica* can develop cross-resistance to several antibiotics under antibiotic-free nutritional stress over hundreds of generations (Knöppel et al., 2017). Oxidative stress has been linked to both ARG emergence (Qi et al., 2023) and transfer (Jong et al., 2020). We reported that ARG abundance in biological wastewater treatment correlates with oxidative-stress response genes, including *oxyR, soxR, sodAB, katG*, and *ahpCF* (Manna et al., 2025). That association was measured at full scale in a system carrying antibiotic residues, where selection on resistance genes cannot be separated from selection on the organisms carrying them. At the biochemical level, respiratory flavoproteins such as fumarate reductase (Frd) and succinate dehydrogenase (Sdh) are major endogenous sources of reactive oxygen species (ROS), generating superoxide (O_2_^•⁻^) and hydrogen peroxide (H_2_O_2_) through the autoxidation of their flavin adenine dinucleotide (FAD) cofactors at rates that scale with enzyme expression (Imlay, 2013; Messner and Imlay, 2002, 1999). ROS-induced DNA damage, in turn, activates transcription-coupled nucleotide excision repair (TC-NER), in which the translocase Mfd, acting with RNA polymerase and UvrA, can lead to mutation through the recruitment of translesion polymerases. This role has led to Mfd being described as an evolvability factor (Carvajal-Garcia et al., 2023; Ragheb et al., 2019). Together, these results outline a chain from physiology to mutation, in which active metabolism generates ROS, ROS damages DNA, and the ensuing repair can itself be mutagenic.

Yet this chain has been assembled almost entirely in single strains under defined laboratory stress, not in complex assemblages responding to fluctuating conditions (Chen et al., 2017; Li et al., 2023). The relationship between the frequency of redox fluctuation, community-wide ROS generation, and the resistome has not been characterized. This gap is significant because seasonal fresh-water systems cycle repeatedly between oxic and anoxic states, creating distinct biogeochemical zones (Diao et al., 2017), and because the timescale of that cycling, whether several times per bacterial generation or over much longer periods, should impose different pressures on microbial metabolism (Chen et al., 2017; Li et al., 2023). Further, many wastewater treatment plants utilize oxic and anoxic zones for nutrient removal. Whether and how this frequency dependence reshapes resistance without antibiotic selection has not been tested.

We hypothesize that the frequency of redox fluctuation is a driver of resistance in freshwater microbial communities. Li *et al*. showed that fast cycling selects for generalists co-expressing both respiratory modes (Li et al., 2023); we reason that this continuous respiration keeps ROS-generating flavoproteins persistently active, thereby imposing chronic oxidative stress (Zhou et al., 2025). We therefore expect communities under high-frequency cycling to show elevated oxidative-stress response transcription, induction of the transcription-coupled repair machinery, and a restructured resistome, relative to communities under slower cycling. We further hypothesize that transcription-coupled repair preferentially targets highly transcribed loci, concentrating any resulting mutational signals on the genes most significantly upregulated by this regime, specifically within the respiratory and oxidative-stress systems, rather than distributing mutations evenly across the genome. To test these expectations, we integrate metagenomics, metatranscriptomics, and strain-resolved population genomics of a controlled redox-cycling freshwater community (Li et al., 2023), characterizing how redox frequency links community-wide ROS and DNA-repair activity to the genomic selection acting on the dominant lineage and to the state of the resistome.

## 2. Methods

### 2.1 Omics Data Collection

For this study, we analyzed publicly available metagenomics and metatranscriptomics datasets generated by Li *et al*. from their controlled chemostat experiments investigating microbial responses to redox fluctuations (Li et al., 2023). The study subjected sulfidic stream microbial communities to alternating oxic-anoxic conditions under three frequency treatments: high frequency (HF, 0.5-day phases), medium frequency (MF, 2-day phases), and low frequency (LF, 8-day phases). The experimental design maintained identical dilution rates (0.5 volume changes per day) across all treatments. Samples were collected at the end of the final oxic and anoxic phases from each triplicate chemostat treatment. For our analysis, we considered samples from all three frequency treatments (high, medium, and low), resulting in six samples (n=6) per condition (three biological replicates × two redox phases). Both phase samples within a treatment experienced the same cycling regime, and the redox phase at sampling was not the variable under test; therefore, the two phases were treated as replicate observations of the treatment. Metagenomic sequencing was performed on a NextSeq 500 system (Illumina), while metatranscriptomic libraries were generated on a MiSeq system (Illumina). We accessed these datasets from NCBI BioProject PRJNA749639 and applied our analytical framework, which focused on oxidative stress metabolism and antibiotic resistance gene dynamics, to address research questions not explored in the original publication.

### 2.2 Metagenomics and Metatranscriptomics Analysis

We performed preprocessing steps to eliminate low-quality sequences using Trimmomatic v0.39 (Bolger et al., 2014) with specific quality thresholds (Manna et al., 2025). For metagenomic analysis, we employed the SqueezeMeta v1.6.0 pipeline (Puente-Sánchez et al., 2020; Tamames and Puente-Sánchez, 2019). Assembly was performed using Megahit v1.2.9 (Li et al., 2015), followed by the removal of redundant contigs using cd-hit (Schmieder and Edwards, 2011a) and merging with Minimus2 (Treangen et al., 2011). Short contigs (<200 bp) were filtered out using prinseq (Schmieder and Edwards, 2011b). The metagenomes and metatranscriptomes were assembled separately and then merged into a single dataset using SqueezeMeta’s merge functionality. RNA sequences were predicted using Barrnap v0.9 (Seemann, 2014), with 16S rRNA sequences taxonomically classified using the RDP classifier (Wang et al., 2007). tRNA and mRNA sequences were identified using Aragorn (Laslett and Canback, 2004), while open reading frames (ORFs) were predicted using Prodigal v2.6.3 (Hyatt et al., 2010). Taxonomic ranks were assigned against the NCBI GenBank nr database with identity thresholds of 85, 60, 55, 50, 46, 42, and 40% for species, genus, family, order, class, phylum, and superkingdom ranks, respectively. Functional annotation was performed through similarity searches using Diamond v2.0.14 (Buchfink et al., 2014) against multiple databases, including GenBank (Clark et al., 2016) and KEGG (Kanehisa and Goto, 2000). Read mapping against assembled contigs was performed using Bowtie2 (Langmead and Salzberg, 2012) and Bedtools (Quinlan and Hall, 2010).

Antibiotic resistance genes were identified from the assembled community open reading frames using two curated tools: the Comprehensive Antibiotic Resistance Database (CARD) Resistance Gene Identifier (RGI) v6.0.5 with CARD v4.0.1 (Alcock et al., 2020), retaining Strict hits, and AMRFinderPlus v4.0.23 (Feldgarden et al., 2021). The union of both tool sets was taken, and open reading frames sharing a gene name were summed to gene level. Gene abundance (metagenome) and expression (metatranscriptome) were quantified as transcripts per million (TPM), normalized by gene length and library size, enabling direct comparison of genetic potential and expression for the same genes.

### 2.3 Strain-resolved population genomic analysis

To resolve within- and between-population variation in the dominant lineage, the quality-filtered reads from all 18 samples were mapped with Bowtie2 (Langmead and Salzberg, 2012) to a concatenated reference of three *Pseudomonas* metagenome-assembled genomes (Li et al., 2023) (MAG A, GenBank GCA_019459105.1; MAG B, GCA_019459245.1; MAG C, GCA_019459405.1), and open reading frames on the reference were predicted with Prodigal v2.6.3 (Hyatt et al., 2010). Each sample was profiled with inStrain v1.10.0 (Olm et al., 2021) using inStrain profile with the reference gene predictions and a scaffold-to-bin file, in --database_mode (minimum read-pair identity 0.92), computing per-genome coverage, breadth, nucleotide diversity and the per-gene ratio of nonsynonymous to synonymous nucleotide variation (pN/pS); variants were called at the inStrain defaults (5× coverage, 0.05 frequency). Between-sample similarity was assessed with inStrain compare, yielding consensus and population single-nucleotide differences and the corresponding average nucleotide identities (conANI, popANI). Genome-wide pN/pS was summarized per sample as the pooled ratio of nonsynonymous to synonymous variant counts across genes passing quality control (coverage ≥ 5, breadth ≥ 0.5). For the gene- and pathway-level comparison, genes meeting the same thresholds in at least four of six samples in both the high- and low-frequency groups were retained (Section 2.4). Analyses focused on MAG A, the lineage that dominated the community; MAG B and MAG C were excluded owing to insufficient or inconsistent coverage across samples.

### 2.4 Statistical Analysis and Visualization

All analyses and visualizations used R version 4.5.1 (Team, 2023), with abundance and expression expressed as transcripts per million (TPM). Heatmaps of ARG genetic potential and expression (Figure 1) display log_2_(TPM + 1), whereas the flavoprotein and electron-transport heatmap (Figure S3) displays per-row z-scores of TPM. Per-gene expression across the three treatments was assessed by Kruskal-Wallis test with Dunn post-hoc contrasts, and total ARG abundance and expression by Kruskal-Wallis test. Genome-level inStrain metrics (coverage, breadth, nucleotide diversity, SNV and SNS density, and genome-wide pN/pS) were compared the same way, and treatments sharing a compact-letter label did not differ significantly. Consensus SNPs between treatment combinations (Figure 4c) were assessed by Wilcoxon rank-sum test; because these are pairwise values from the same six samples per treatment, the comparison is indicative of the divergence distributions and not a test on independent replicates. For gene- and pathway-level selection, per-gene log_2_ fold changes in median pN/pS (HF versus LF) were tested by Wilcoxon rank-sum test and aggregated to KEGG level-3 pathways by one-sample Wilcoxon test against zero, with pathways ranked and displayed by effect size at a cutoff of |median log_2_FC| > 1. Coverage and pN/pS were related by Spearman rank correlation. Sankey diagrams of ARG taxonomic contributors, based on contig assignment, were built with networkD3. Kruskal-Wallis p-values are unadjusted where one test was run per metric and Benjamini-Hochberg adjusted across genes where run over a gene set; Dunn p-values are adjusted across all pairwise contrasts within each gene set, and Wilcoxon and pathway-level p-values are adjusted likewise. Significance was set at p < 0.05.

**Figure 1.**
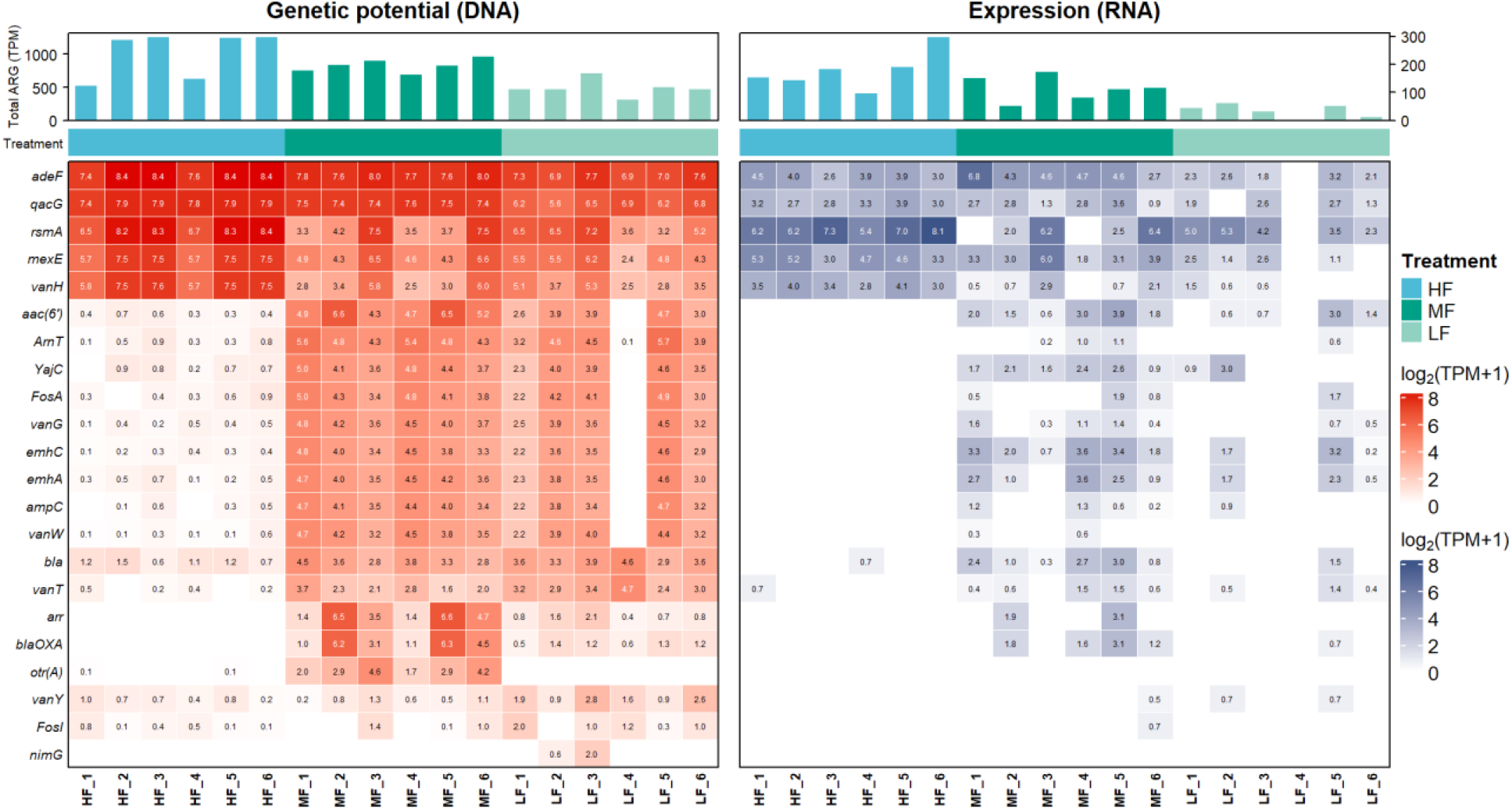
Genetic potential and expression of community ARGs across redox oscillation frequencies. Paired heatmaps of metagenomic abundance (left, genetic potential) and metatranscriptomic expression (right) for 22 ARGs under high-frequency (HF), medium-frequency (MF), and low-frequency (LF) oxic–anoxic cycling (n = 6 per treatment: three replicates × oxic and anoxic phases). Cell values are log_2_(TPM + 1) of DNA and RNA; bar plots above each panel give total ARG TPM per sample. Rows are ordered by hierarchical clustering of the DNA profiles and applied to both panels.

## 3. Results and Discussion

### 3.1 Frequency-dependent expression of ARGs by redox dynamics

To understand how the community resistome responds to redox oscillation frequency, ARGs were identified from the assembled community open reading frames using two curated tools, CARD-RGI (Strict hits) and AMRFinderPlus, and their metagenomic abundance (genetic potential) was compared with their metatranscriptomic gene expression across high-frequency (HF), medium-frequency (MF), and low-frequency (LF) oxic–anoxic cycling (Figure 1). The retained set was dominated by multidrug efflux determinants, including the RND pumps *adeF* and *mexE*, the small multidrug-resistance pump *qacG*, and the efflux regulator *rsmA*, together with a smaller group of enzymatic determinants (β-lactamase-family genes, the aminoglycoside acetyltransferase *aac(6′)*, the rifamycin ARR family, and fosfomycin thiol transferases). Two of the retained determinants are regulatory or accessory rather than resistance genes in the strict sense. The *rsmA* is a CsrA-family post-transcriptional regulator, curated in CARD for its control of efflux expression, and *vanH* is a dehydrogenase of the vancomycin gene cluster whose assignment in a *Pseudomonas* background should be treated as provisional. Both are retained as part of the curated tool output.

At the level of genetic potential, efflux genes were present in all treatments. Their metagenomic abundance was highest under HF (summed ARG abundance, median per sample HF = 1227, MF = 833, LF = 472 TPM; p = 0.0071), but the genes themselves were detectable across HF, MF, and LF. The enzymatic determinants (the aminoglycoside acetyltransferase *aac(6′)*, the β-lactamases *blaOXA* and *ampC*, the rifampin ADP-ribosyltransferase *arr*, and the fosfomycin enzyme FosA) showed a different distribution. They were largely confined to the MF and LF metagenomes and were sparse or absent under HF (*aac(6′)*: HF 0.4, MF 48.9, LF 10.8 TPM; *blaOXA*: HF 0.0, MF 29.8, LF 1.1 TPM). Genetic potential for resistance was therefore broad but unevenly distributed, with the widest gene content under MF and LF.

Expression, by contrast, followed a more structured pattern. Total ARG transcription was highest under HF and lowest under LF (median per sample HF = 168, MF = 113, LF = 38 TPM; p = 0.0027), but this total was carried by a small number of efflux genes. The efflux regulator *rsmA* and the RND pump *mexE* reached their highest expression under HF (*rsmA*: p = 0.014, q = 0.030; *mexE*: p = 0.0047, q = 0.016), whereas *adeF*, the most prevalent efflux gene (p = 0.0034, q = 0.016), and nearly all of the enzymatic determinants (*aac(6′)* q = 0.016; *bla* q = 0.016; *blaOXA* q = 0.036) reached their highest expression under MF. The number of distinct ARGs expressed per sample was roughly 2-fold higher under MF than under HF or LF. Redox frequency therefore reshapes which resistance genes are expressed. High-frequency cycling concentrates transcription in a few efflux genes, and medium-frequency cycling supports a broader, more even resistome.

This restructuring occurred with no antibiotic added to the system (Andersson and Hughes, 2012; Knöppel et al., 2017). The medium was chemically defined and antibiotic-free (Li et al., 2023), and continuous dilution removed any inoculum-borne residue; because the same inoculum and medium were split across the three frequencies, no residual compound could produce a frequency-dependent effect. The result establishes redox oscillation frequency as an environmental variable that sets the transcriptional state of the resistome, and it separates two properties that resistance surveillance usually treats as one: the resistance genes a community carries, and the genes it actually expresses.

The expressed resistome concentrated in a single lineage (Figure 2). Under HF, 99% of ARG transcription originated from *Pseudomonas*, across only 4 active taxa. MF was the most diverse case, 70% *Pseudomonas* across 8 taxa, with about a third of its resistome expression contributed by other *Proteobacteria* (Figure S1). LF spread across 11 taxa and remained *Pseudomonas*-leaning (86%) but transcriptionally quiet (Figure S2), carrying the lowest total expression of the three treatments (1059, 682, and 201 TPM summed across the six replicates of HF, MF, and LF, respectively). The concentrated HF resistome therefore tracks the composition of the active community rather than gene-specific regulation: it is one lineage’s constitutive efflux program (*rsmA, mexE, qacG*), expressed in proportion to how completely that lineage dominates the transcription. This dominance reflects the lineage itself: *Pseudomonas* carries an unusually large set of chromosomally encoded RND efflux systems, including MexEF-OprN and its regulator

**Figure 2.**
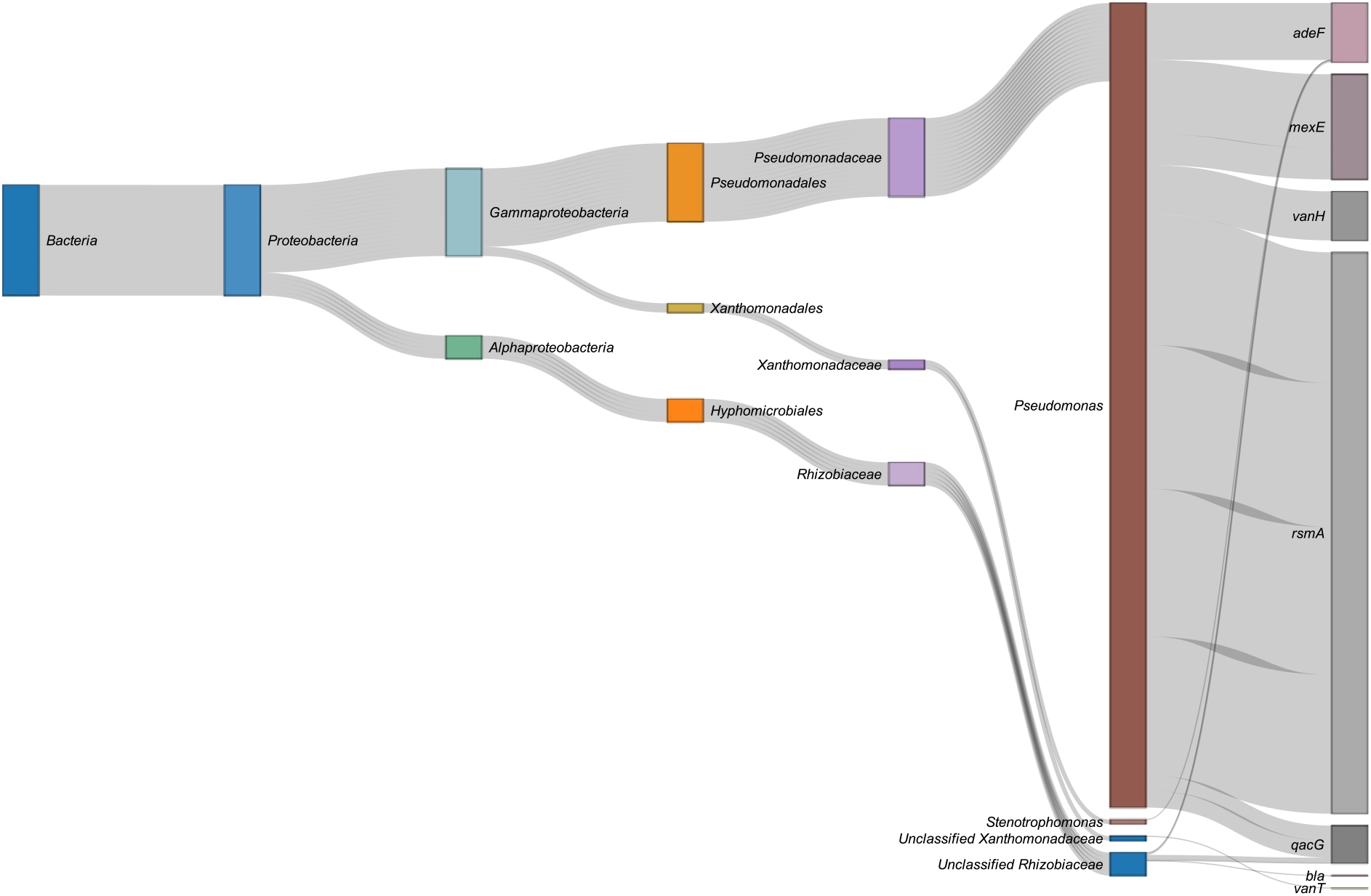
Hosts of ARGs expressed under high-frequency (HF) redox cycling. Sankey diagram of ARG transcript flow across taxonomic ranks (from kingdom to genus), with hosts assigned based on contig taxonomy. Edge width between taxonomic ranks is the number of gene occurrences; genus-to-gene edge width is proportional to mean per-gene TPM. Most transcripts originated from *Proteobacteria*, chiefly *Gammaproteobacteria* and *Alphaproteobacteria*, and were dominated by the genus *Pseudomonas*.

RsmA, as an intrinsic, constitutive resistome whose transcription scales with the lineage’s activity rather than with antibiotic exposure. Its facultative and denitrifying metabolism also suits rapid oxic-anoxic switching, so high-frequency cycling concentrated the active community, and with it the expressed resistome, onto this single versatile taxon.

### 3.2 Endogenous oxidative stress under high-frequency cycling

The elevated resistome expression under fast cycling occurred with no antibiotic present, which points to the metabolic state as the driver. We examined that state directly, focusing on the enzymes that generate endogenous ROS. Flavoprotein respiratory enzymes are the principal intracellular sources of superoxide and hydrogen peroxide, produced by univalent electron leak to molecular oxygen at their flavin (FAD or FMN) cofactors (Imlay, 2013; Messner and Imlay, 1999), and each returning to oxic conditions produces a burst of superoxide from a reduced respiratory chain, so repeated transitions impose recurrent oxidative load (Kargalioglu and Imlay, 1994). Under fast cycling the community is dominated by generalists that co-express aerobic and anaerobic respiration continuously, rather than shutting down aerobic metabolism during the anoxic phase (Li et al., 2023), so respiratory electron flux, and the leak that accompanies it, persists across both phases.

The core respiratory flavoproteins were induced under HF cycling (Figure S3). Succinate dehydrogenase subunit B (*sdhB*), a Complex II flavoprotein and a documented superoxide source (Messner and Imlay, 2002), rose about 9-fold in HF relative to LF cycling (HF mean 1037 vs LF 119 TPM), and the remaining Sdh subunits followed the same direction (*sdhA* 6.6-fold, *sdhC* 6.7-fold, *sdhD* 7.3-fold). The NADH dehydrogenase (*nuo*) operon did not differ significantly (adjusted p ≥ 0.057 for all 14 subunits) across frequencies and peaked under MF rather than HF, which places the Complex II flavoprotein, not respiratory chain expression in general, at the centre of the signal. Terminal electron transfer increased in parallel, with the cytochrome bc_1_ complex elevated about 6-fold (*petA*), 8-fold (*petB*) and 9-fold (*petC*), and cytochrome c (CYC) reaching about 15-fold over LF (HF 1806 vs LF 123 TPM). The high-affinity *cbb*_*3*_-type cytochrome c oxidases, whose low oxygen K_m_ supports respiration under microaerobic conditions (Arai et al., 2014), rose in parallel across all three subunits (*ccoN* 6.2-fold, *ccoO* 6.3-fold, *ccoP* 4.5-fold), consistent with a community respiring through both the oxic and the low-oxygen phases of each cycle. Citrate synthase (*gltA*), a non-flavoprotein marker of TCA-cycle entry, tracked the same pattern (about 5-fold; HF 700 vs LF 148 TPM), which places the signal at carbon flux into the cycle rather than at a single enzyme. Together, these fold changes place the community in a high-flux aerobic respiratory state, the condition under which electron leak and ROS output are highest.

The antioxidant response moved with the ROS sources. Glutathione reductase (*gor*), which regenerates reduced glutathione during oxidative stress, was about 4-fold higher in HF (HF 72 vs LF 17 TPM), and lipoamide dehydrogenase (*lpd*), the flavoprotein shared across the 2-oxoacid dehydrogenase complexes, rose about 12-fold (HF 748 vs LF 64 TPM). Co-induction of oxidant sources and thiol-based defenses is the transcriptional signature expected when a cell is managing a sustained oxidative burden (Imlay, 2013). The signal extended beyond the respiratory chain. The glutamate synthase subunits *gltB* and *gltD*, which carry FAD/FMN and consume NAD(P)H during nitrogen assimilation, were each about 4-fold higher in HF, and L-aspartate oxidase (*nadB*), an FAD enzyme of NAD biosynthesis (Messner and Imlay, 2002), rose about 5-fold (HF 52 vs LF 10 TPM). These indicate that HF cycling raises redox-reactive flux across nitrogen and cofactor metabolism, not only in core respiration. All genes named above differed significantly across cycling frequencies (Kruskal-Wallis with Benjamini-Hochberg correction, adjusted p ≤ 0.025).

The respiratory induction, the parallel antioxidant response, and the extension into nitrogen and cofactor metabolism together describe cells generating endogenous ROS under repeated oxygen transitions and buffering it (Figure S3). This transcription was confined to *Proteobacteria* and concentrated in *Pseudomonas* under HF (69% of electron transport chain gene expression), whereas under MF it spread across more genera and the *Pseudomonas* share fell to 28% (LF 51%; Figures S4–S6). The ROS-generating respiratory state is therefore the metabolic condition that fast cycling imposes on the dominant lineage.

### 3.3 Antioxidant defense response across redox cycling frequencies

The respiratory flavoproteins that generate endogenous ROS create a matching demand for its removal. We examined the expression of dedicated antioxidant defense genes across the frequency gradient (Figure 3). Most were coordinately induced under HF cycling relative to LF cycling, though the response was uneven and several changes carried high between-replicate variance. For example, the peroxide-responsive regulator *oxyR* (Imlay, 2013) rose about 6-fold in HF (HF mean 58 vs LF 10 TPM; adjusted p = 0.007), whereas the superoxide-responsive *soxR* increased about 2.3-fold, a change that was not significant (HF 18 vs LF 8 TPM; adjusted p = 0.11). The *oxyR* induction points to H_2_O_2_ as the dominant sensed stress under HF. The single largest change was *cybB*, encoding a membrane cytochrome b, which rose about 58-fold (HF 1566 vs LF 27 TPM). The magnitude is large, and the HF replicates are dispersed, though the rank-based test is insensitive to that dispersion (adjusted p = 0.006). *CybB* may participate in transmembrane superoxide-to-quinone electron transfer, potentially helping buffer periplasmic redox stress (Lundgren et al., 2018). Among the peroxide-degrading enzymes, catalase *katE* rose about 7-fold (HF 171 vs LF 25 TPM; adjusted p = 0.016). The catalase-peroxidase *katG* changed by a larger amount, about 9-fold (HF 423 vs LF 46 TPM; adjusted p = 0.016), despite high variance across the HF replicates. Superoxide dismutase (*sodAB*) was induced about 6-fold (HF 801 vs LF 141 TPM; adjusted p = 0.006), consistent with a raised superoxide burden under HF.

**Figure 3.**
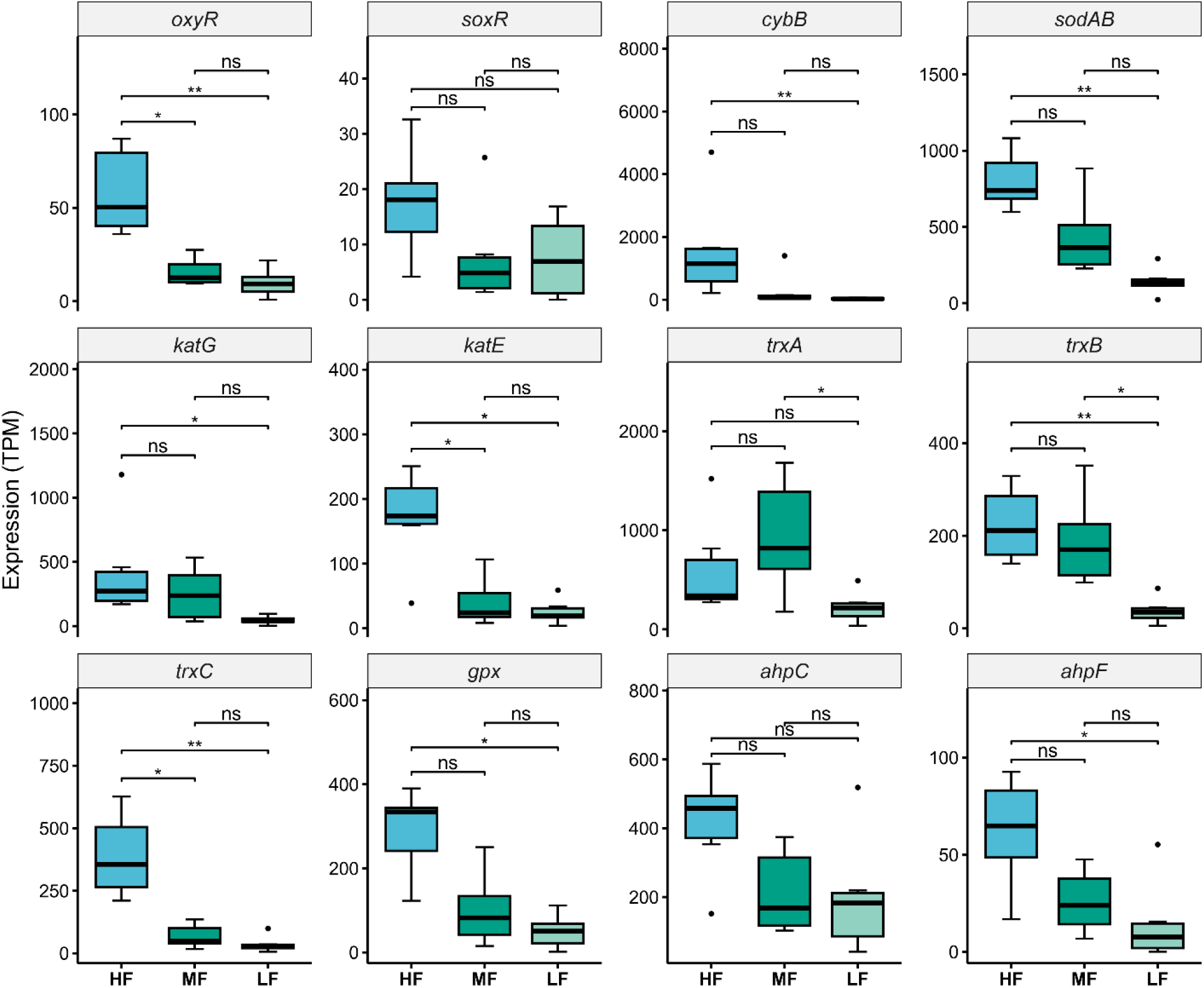
Expression of oxidative-stress response genes across redox fluctuation frequencies. Boxplots of transcript abundance (TPM) for oxidative-stress response genes under high- (HF), medium- (MF), and low-frequency (LF) oxic–anoxic cycling (n = 6 per treatment). Panels are ordered by function: regulators (*oxyR, soxR*), cytochrome b (*cybB*), ROS scavengers (*sodAB, katG, katE*), the thioredoxin system (*trxA, trxB, trxC*), glutathione peroxidase (*gpx*), and alkyl hydroperoxide reductase (*ahpC, ahpF*). Significance was assessed by Kruskal-Wallis tests with Dunn post-hoc contrasts, both corrected by the Benjamini-Hochberg procedure: ns, not significant; * adjusted p < 0.05; ** adjusted p < 0.01.

The thiol-based redox-buffering systems were the most consistently induced. Thioredoxin *trxC* rose about 11-fold (HF 389 vs LF 36 TPM; adjusted p = 0.006) and thioredoxin reductase *trxB* about 6-fold (HF 224 vs LF 38 TPM; adjusted p = 0.008). Glutathione peroxidase (*gpx*) increased about 6-fold (HF 290 vs LF 50 TPM; adjusted p = 0.012). The alkyl hydroperoxide reductase subunit *ahpC* increased about 2-fold, but this change was not significant (HF 417 vs LF 200 TPM; adjusted p = 0.11), whereas its partner subunit *ahpF* rose about 4-fold (HF 62 vs LF 15 TPM; adjusted p = 0.017). These systems reduce oxidized protein thiols and detoxify organic peroxides.

Taken together, HF cycling drove a coordinated antioxidant response spanning the peroxide and superoxide arms, as well as both the catalase and thiol pathways (Figure 3). The strongest and most reproducible signals were in the thioredoxin system, *sodAB*, and *oxyR*, with both catalases (*katE, katG*) and *ahpF* also significant. The breadth and magnitude of the significant responses indicate sustained intracellular oxidative pressure during rapid redox transitions, consistent with ROS-generating respiratory and flavoprotein metabolism. As with respiration, this antioxidant response was *Proteobacterial* and concentrated in *Pseudomonas* under HF (78% of antioxidant gene expression), falling to 28% under MF as the community broadened (LF 52%; Figures S7– S9). The HF antioxidant response, therefore, sits in the same dominant Proteobacterial lineage as the respiratory signal, so the oxidative pressure and its buffering fall on the same organisms.

### 3.4 Induction of transcription-coupled DNA repair under high-frequency cycling

A cell sustaining high-flux aerobic respiration under repeated oxygen transitions incurs oxidative damage to DNA. Reactive oxygen species oxidize bases and can stall replication and transcription, and cells respond by inducing nucleotide excision repair (NER) and translesion synthesis (Carvajal-Garcia et al., 2023; Ragheb et al., 2019). We examined whether this response was detectable in the metatranscriptome and found it was induced specifically under HF cycling (Figure S10). The NER pathway was induced coordinately under HF. The excision nuclease subunits *uvrA, uvrB*, and *uvrC* rose 7.4-, 12.7-, and 6.3-fold over LF (*uvrA* HF 92 vs LF 12 TPM, adjusted p = 0.003; *uvrB* HF 88 vs LF 7 TPM, adjusted p = 0.007; *uvrC* HF 25 vs LF 4 TPM, adjusted p = 0.003). The coupling factor *mfd*, which removes stalled RNA polymerase from the template and recruits UvrA to lesions on the transcribed strand (Selby and Sancar, 1993), rose 5.8-fold (HF 36 vs LF 6 TPM, adjusted p = 0.003), and the RNA polymerase subunits *rpoC* and *rpoB* were elevated in parallel (*rpoC* 6.5-fold, HF 726 vs LF 111 TPM, adjusted p = 0.012). This pattern is characteristic of transcription-coupled repair rather than general genome maintenance. The error-prone polymerases were induced in parallel. *dinB* (polymerase IV) and *umuC* (the catalytic subunit of polymerase V), the two Y-family translesion enzymes, rose 5.4- and 6.4-fold (*dinB* HF 15 vs LF 3 TPM, adjusted p = 0.009; *umuC* HF 28 vs LF 4 TPM). The high-fidelity gap-filling polymerase *polA* rose 5.1-fold (HF 42 vs LF 8 TPM, adjusted p = 0.003). For *umuC*, expression differed across the three frequencies (Kruskal-Wallis adjusted p = 0.044), but the HF versus LF contrast alone did not reach significance (adjusted p = 0.056), so its induction is suggestive rather than established. Coordinated induction of excision repair and, in dinB, of error-prone synthesis is the transcriptional profile expected of cells clearing oxidative lesions under load, and it is the configuration through which repair can itself introduce mutations.

A consequence of transcription-coupled repair is preferential mutation accumulation in highly transcribed genes (Ragheb et al., 2019), although mutation rates were not directly measured here. Under HF, the most highly transcribed genes are those involved in redox and central metabolism, not resistance genes, so the mutational bias falls on metabolic loci. Transcription-coupled repair, therefore, acts on metabolism rather than on the resistome, and this separation is the decoupling at the center of the study. The repair response was concentrated in one part of the community. We traced the taxonomic origin of the HF repair transcripts (Figure S11): all originated from Proteobacteria, with Pseudomonas the largest source at 61% of repair-gene transcription, followed by *Rhizobiaceae* (21%) and other *Proteobacteria* (19%). Under MF, the *Pseudomonas* share of repair transcription fell to 17% and under LF to 43% (Figures S12–S13), so the repair signal, like the metabolic and antioxidant ones, concentrates the mutational pressure on the eventual dominant lineage. Across the four transcriptional layers, respiration, antioxidant defense, DNA repair, and the resistome, high-frequency cycling concentrated expression in *Pseudomonas*, and that concentration relaxed as the community broadened under medium and low frequency (Figure S14). Resolving which lineage comes to dominate, and what that dominance does to the resistome, requires strain-resolved genomics of the dominant population.

### 3.5 Redox frequency sets which lineage monopolizes the community

The metatranscriptomic analysis describes what the community expresses but not how redox frequency reorganizes the underlying populations. Because *Pseudomonas* accounted for the largest share of transcription across all four functional layers (Figure S14), we resolved the population structure of this genus directly using strain-resolved metagenomics (inStrain). *Li et al*. recovered three *Pseudomonas* metagenome-assembled genomes (MAGs) from this system, MAGs A, B, and C (Li et al., 2023). Only MAG A was recovered at near-complete breadth of coverage at 5× or more (median 99.9% under HF, 98.8% under MF and 99.5% under LF) in all three treatments, giving the coverage that a per-gene selection analysis requires (Figure S15).

MAG B was absent from all high-frequency replicates, so it cannot be compared across the gradient. MAG C was only partially recovered under HF (median 66% of the genome at 5× or more) and MF (17% at 2.6× coverage), too sparse for genome-wide statistics. We therefore focused on MAG A, the dominant *Pseudomonas* genome (Figure 4). *Li et al*. showed that fast cycling selects a single generalist community at the population level (Li et al., 2023); we ask how that selection acts within the dominant lineage. One variable organizes the patterns that follow, the degree to which a single metabolic-generalist lineage dominates the community, set by cycling frequency (Chen et al., 2017; Diao et al., 2017). Fast cycling drove a clonal sweep. Under HF, MAG A reached a median coverage of 331× against 24× under LF (Kruskal-Wallis p = 0.01, Dunn HF–LF adjusted p = 0.01; Figure 4a). HF replicates converged on a shared consensus, differing by a median of 173 consensus SNPs, whereas LF replicates differed by 1442 (Figure 4c). Because a consensus difference is called wherever the majority allele differs, these counts track the stability of the majority call, not allele content, which was effectively identical, since population ANI reached 1.000000 in every treatment with one to two population single-nucleotide differences across the 4.3 to 4.6 Mb compared. MF replicates were at least as tightly converged as HF (151 consensus SNPs, conANI 0.999966; Figure 4d), so this deterministic selection was not confined to the fastest regime. Nucleotide diversity did not differ across treatments (median about 0.0027, Kruskal-Wallis p = 0.74; Figure 4e) and SNV density was higher in LF (2.3 vs 0.7 SNVs per kb, Kruskal-Wallis p = 0.28; Figure 4f), a difference not explained by detection power, since HF had the deepest sequencing, and linkage is also lowest under HF (r2 0.74 against 0.92 under LF), as expected when a single genotype dominates and co-inherited variants raise linkage in mixed populations. Coverage is not normalised to library size, so it indexes how completely this lineage was sampled, not its exact share of the community. The HF signature is therefore the deterministic selection of one lineage across replicates, accompanied by a reduction in the variation that lineage carries. This consensus-level convergence does not conflict with the greater between-replicate variability of HF transcriptomes reported previously (Li et al., 2023), since selection acts deterministically on the genome while expression remains stochastic.

**Figure 4.**
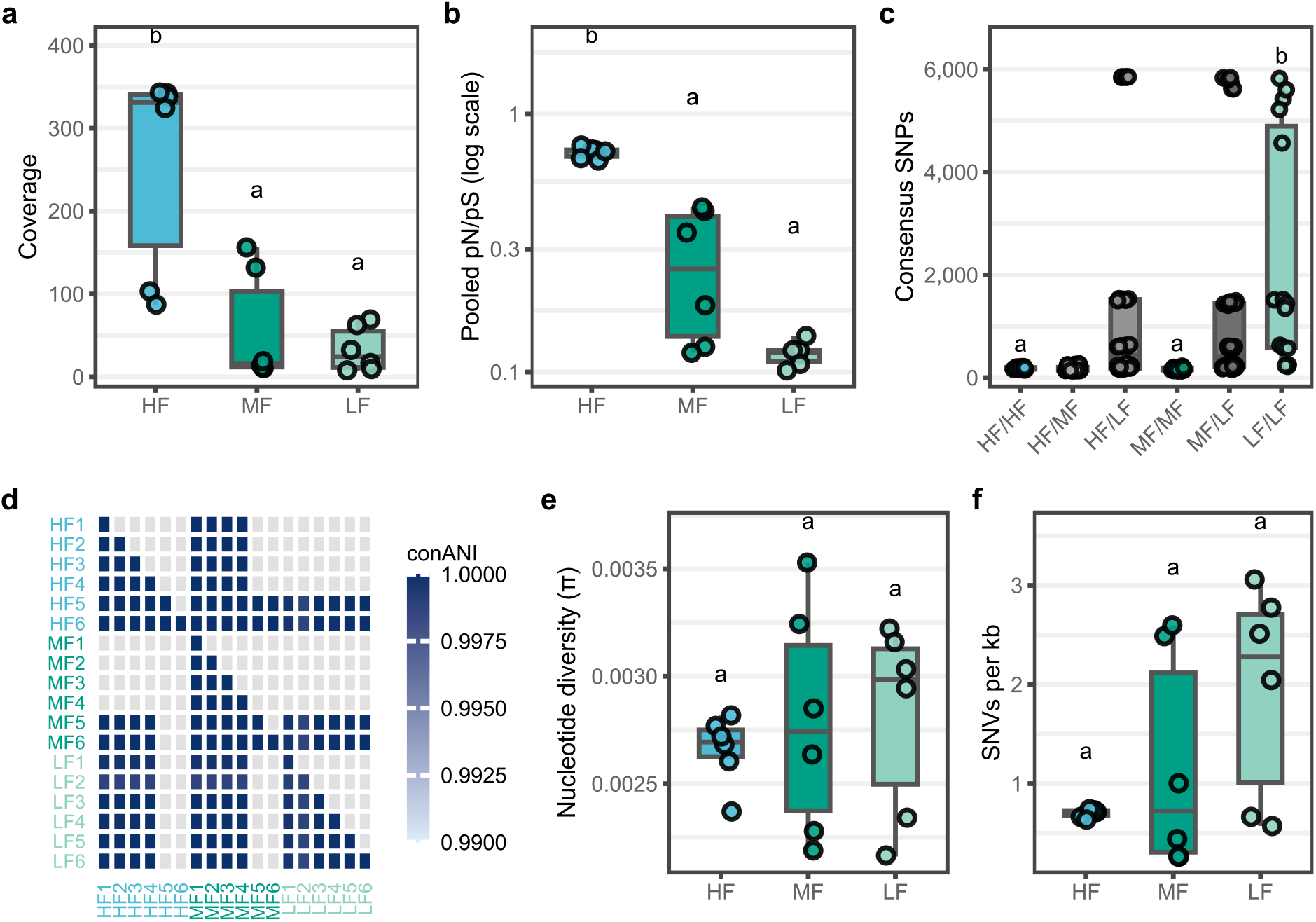
Population genomics of the dominant *Pseudomonas* lineage (MAG A) across redox fluctuation frequencies. Strain-resolved inStrain metrics under high- (HF), medium- (MF), and low-frequency (LF) oxic–anoxic cycling, n = 6 per treatment. Panels show (a) coverage, (b) genome-wide pooled pN/pS (log scale), (c) consensus SNPs for every sample pair by treatment combination, (d) pairwise consensus ANI (conANI) across all 18 samples, (e) nucleotide diversity (π), and (f) SNV density (per kb). Compact letters mark groups with no significant difference (shared letter), Kruskal–Wallis with Dunn in (a, b, e, f), Benjamini–Hochberg corrected; letters in (c) mark an indicative Wilcoxon rank-sum comparison of pairwise consensus-SNP counts. popANI is 1.000000 within every treatment, so the treatments differ in the stability of the majority allele call while their allele content is indistinguishable.

However, the resistance genes did not evolve. The four ARGs in MAG A carried zero SNVs across all six HF replicates, despite HF having the deepest coverage at these loci (188× to 290×), and accumulated variation only under MF (22 SNVs) and LF (52 SNVs), of which 17 of 22 and 49 of 52 were synonymous (Figure S16). None of the four genes carried a fixed difference from the reference in any of the 18 samples, so their consensus sequences were identical throughout. Under HF the dominant genotype had fixed, leaving its ARG alleles as the only ones present, which accounts for the absence of variants without invoking any protection from mutation. The MF and LF variants represent neutral standing diversity among coexisting strains under purifying selection. Resistance sequences did not change over the course of this experiment at any frequency.

Selection acted on MAG A’s metabolic genome, while its resistance genes stayed conserved. Genome-wide, MAG A was under purifying selection in every treatment (pooled pN/pS 0.72 in HF, 0.26 in MF, 0.12 in LF, all below 1). Purifying selection was weakest under HF, where its pN/pS, though still below 1, was higher than under MF or LF, meaning more nonsynonymous variation was tolerated (Kruskal-Wallis p = 0.0011; Dunn HF-LF adjusted p = 0.0007, HF-MF adjusted p = 0.046, MF-LF adjusted p = 0.13; Figure 4b). Coverage and pN/pS correlated across samples (Spearman rho = 0.92), raising the possibility that the higher pN/pS under HF was coverage-driven, a known concern in metagenomic population analyses (Rodriguez-R and Konstantinidis, 2014). We therefore treat the elevated HF ratio as a qualitative signal of relaxed constraint and not a precise estimate. The genes carrying the relaxed-constraint signal trend toward the oxidative-stress and redox-cofactor pathways induced under fast cycling (glutathione, porphyrin, folate, nicotinate and NAD, nitrogen; Figure 5). This pathway signal is exploratory, an effect-size comparison that does not survive multiple-testing correction at six replicates per treatment, so we interpret it as a distributional shift, a modest genome-wide relaxation concentrated on the most-transcribed metabolic genes and not positive selection at individual loci.

**Figure 5.**
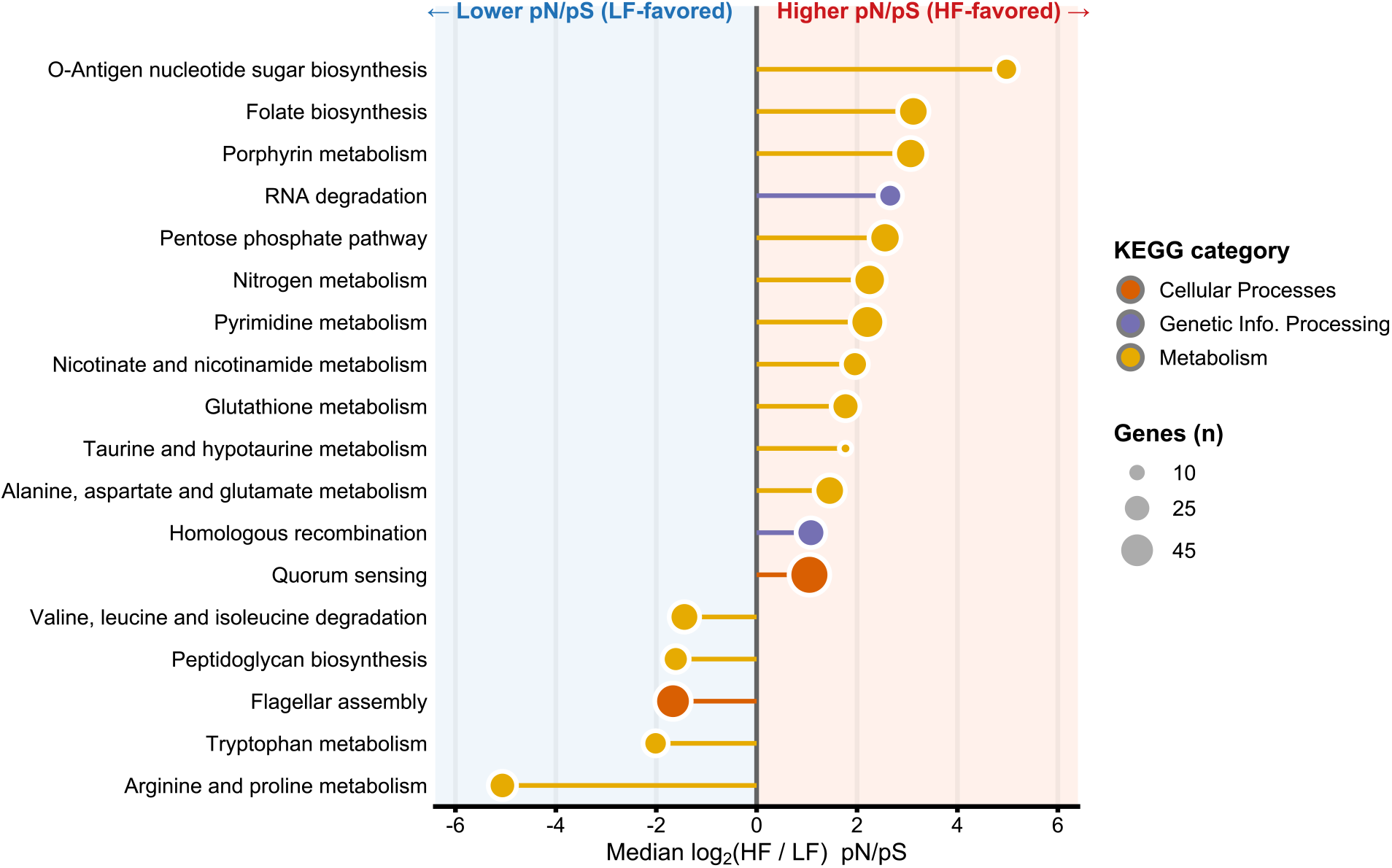
Pathway-level shift in selection (pN/pS) between high- and low-frequency cycling in *Pseudomonas* MAG A. Each point is a KEGG level-3 pathway, plotted as the median log_2_ fold change in per-gene pN/pS between high-frequency (HF) and low-frequency (LF) cycling; positive values (red) indicate relaxed purifying selection under HF. Only pathways with |median log_2_FC| > 1 are shown. Color denotes the KEGG level-1 category, size denotes the number of genes. Pathways are ranked by effect size; no pathway was significant after Benjamini–Hochberg correction at n = 6, so the figure shows a distributional shift in constraint and not selection at named loci.

The resistance genes remained in the purifying background and were sequence-conserved, while their expression increased. Redox frequency amplifies resistance by selecting and monopolizing a high-efflux metabolic generalist, and that lineage co-regulates efflux with its metabolic state. Resistance therefore rises as a consequence of metabolic selection on which lineage dominates. The ROS and repair signatures are genuine, but they act on the metabolism of the dominant lineage, and resistance-gene expression rises with it.

### 3.6 Limitations

Several constraints bound these conclusions. The data are a reanalysis of a single controlled chemostat experiment from one published dataset, so the findings need confirmation in independent systems before they generalize. There is no time-zero sample, so the comparison is cross-sectional across redox regimes and cannot capture gains or losses in resistance over time. We did not measure reactive oxygen species, so the oxidative-stress step in the proposed mechanism is inferred from redox condition and repair-gene induction rather than observed. Resistance is inferred from transcript abundance, not from a measured resistance phenotype, so the decoupling we report is between sequence-level selection and expression, not between sequence and a functional assay. The transcriptional layer (Sections 3.1 to 3.4) is community-wide, whereas the selection analysis is resolved to a single genome (MAG A). The taxonomy panels tie the two together but do not conflate them into a single measurement. The population-genomic conclusions rest on the dominant *Pseudomonas* lineage, because the other two *Pseudomonas* genomes were either absent at high frequency or recovered with insufficient breadth (Figure S15), so responses in less-abundant populations remain unresolved. The transcription-coupled mutational bias is a candidate mechanism, inferred from coordinated repair-gene induction and the genomic location of variation rather than measured directly.

## 4. Conclusion

Fast redox cycling raised antibiotic resistance expression in a sulfidic stream community with no antibiotic present. Surveillance built on which resistance genes are present and how their sequences change does not capture this risk. Under fast oxic-anoxic cycling, and with no antibiotic added, community-wide resistance expression rose, yet the efflux genes carrying that increase were present in every regime and showed no sequence change, carrying no fixed difference from the reference in any sample, so neither their presence nor their sequence evolution signalled the increase. Total gene abundance across the community also rose under fast cycling, but it tracked the dominance of one host lineage rather than any change in the resistome, and the rise in expression exceeded the rise in abundance. What changed was which lineage dominated the community and how hard it expressed a constitutive efflux program. A survey based on the presence or sequence of these efflux genes would score the three regimes as equivalent, and the wider gene inventory would rank medium- and low-frequency cycling as the broader resistomes, inverting the expression ranking. One based on gene abundance would read the rise as more resistance genes, when what rose was how heavily a single lineage expressed a constitutive efflux program. The determinant here is a physical variable, the frequency of redox change, acting through selection on metabolism rather than through antibiotic pressure. Oxygen availability fluctuates in many natural and engineered water systems, including sediments, biofilms, stratified waters, and reactors operated with intermittent aeration, making redox dynamics a candidate control point for where resistance expression is concentrated. Testing whether the taxon-selection-and-expression mechanism seen here operates in situ, and whether expression-aware monitoring adds predictive value over gene-based surveillance, is the practical next step.

## Data Availability

All data analyzed in this study were obtained from publicly available repositories. The multi-omics datasets were originally generated by Li et al. (2023) and are available under Bioproject PRJNA749639 (NCBI), including metagenomic raw reads (Biosamples: SAMN20395938-SAMN20395955), and metatranscriptomic raw reads (Biosamples: SAMN20446884-SAMN20446901).

## Author information

## CRediT authorship contribution statement

**BM**: Conceptualization, Investigation, Methodology, Formal Analysis, Visualization, Writing - original draft. **IK:** Writing – review & editing; **QL:** Writing – review and editing; **NS**: Conceptualization, Resources, Funding acquisition, Writing - review & editing.

## Notes

The authors declare no competing financial interest.

## Acknowledgment

The authors acknowledge the use of New Zealand eScience Infrastructure (NeSI) high-performance computing facilities, consulting support, and/or training services as part of this research. New Zealand’s national facilities are provided by NeSI and funded jointly by NeSI’s collaborator institutions and through the Ministry of Business, Innovation & Employment’s Research Infrastructure program. URL https://www.nesi.org.nz. The authors also acknowledge the Center for eResearch at the University of Auckland for their help in facilitating this research. http://www.eresearch.auckland.ac.nz.

