## Supporting Information for "Redox oscillation frequency reshapes antibiotic resistance-gene expression through metabolic and ecological selection in a freshwater community"

**for**

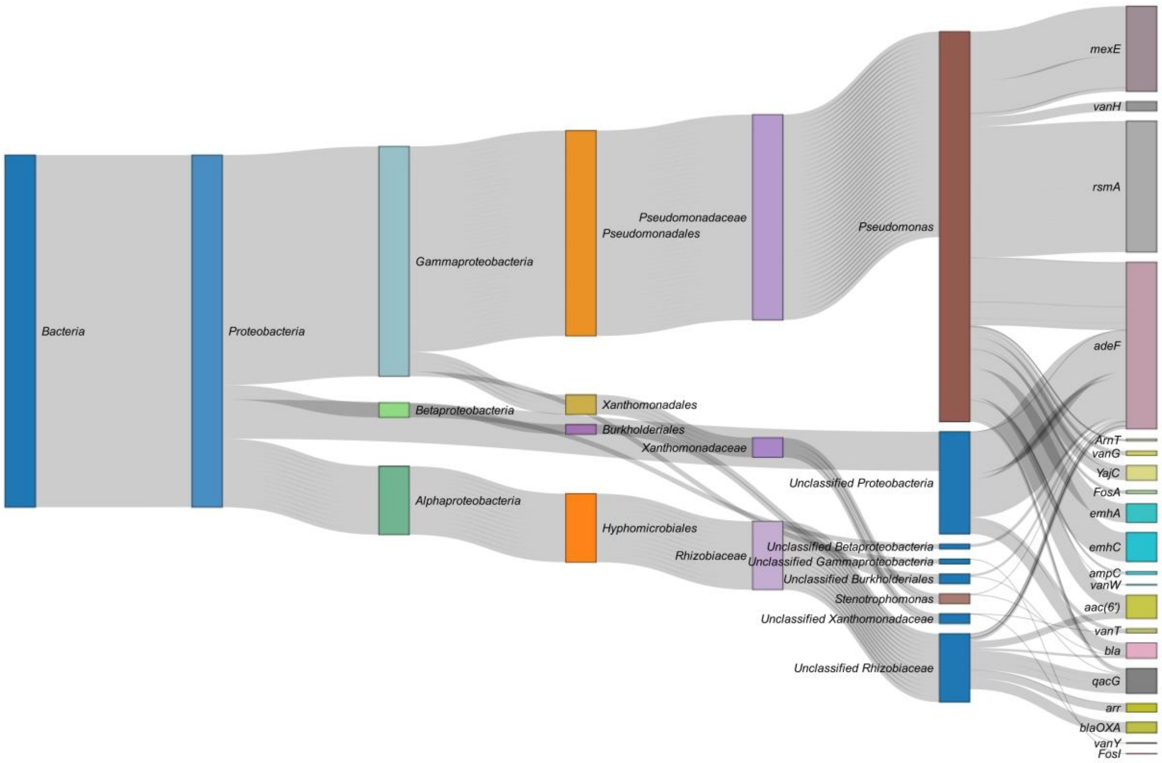


**Figure S1.** Hosts of ARGs expressed under medium-frequency (MF) redox cycling. Sankey diagram of ARG transcript flow across taxonomic ranks (superkingdom to genus), with hosts assigned from contig taxonomy. Edge width between taxonomic ranks is the number of gene occurrences; genus-to-gene edge width is proportional to mean per-gene TPM.

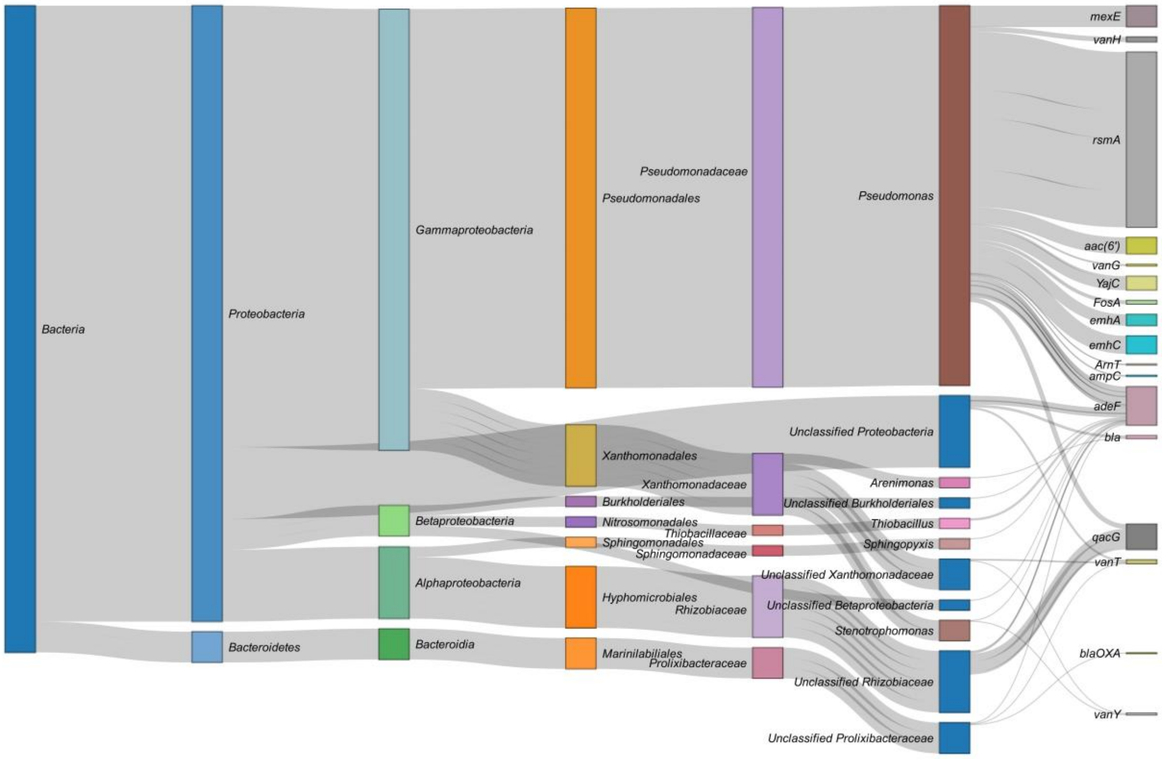


**Figure S2.** Hosts of ARGs expressed under low-frequency (LF) redox cycling. Sankey diagram of ARG transcript flow across taxonomic ranks (superkingdom to genus), with hosts assigned from contig taxonomy. Edge width between taxonomic ranks is the number of gene occurrences; genus-to-gene edge width is proportional to mean per-gene TPM.

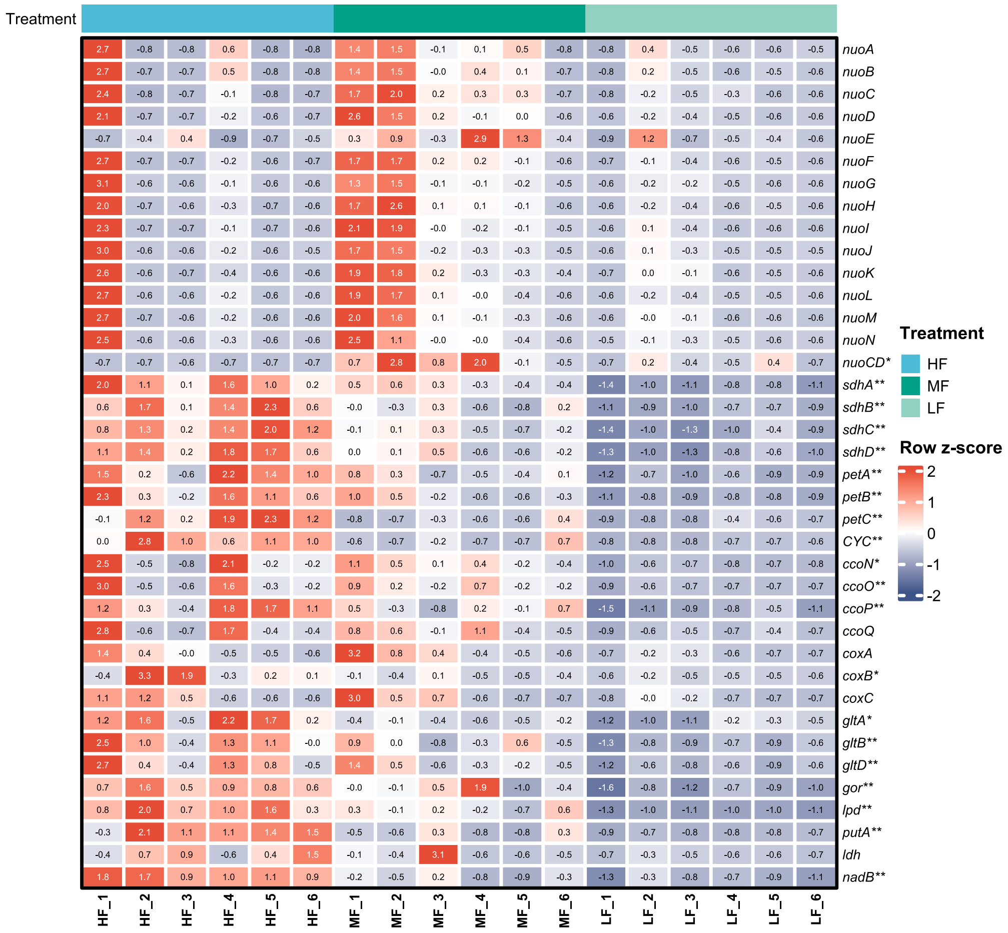


**Figure S3.** Transcriptional responses of flavoprotein oxidoreductases and respiratory enzymes under varying redox fluctuation frequencies. Expression of flavoprotein and electron transport chain (ETC) genes across high-frequency (HF), medium-frequency (MF), and low-frequency (LF) oxic–anoxic cycling (n = 6 per treatment). Cells show the row z-score of transcript abundance (TPM), computed per gene across all 18 samples, with the z-score printed in each cell; genes are shown in pathway order without clustering. Expression of 20 of the 38 genes differed significantly across cycling frequencies. The significant genes span Complex II (*sdhA*–*sdhD*), the bc1 complex (*petA*–*petC*, *CYC*), the cbb3 oxidase (*ccoN*, *ccoO*, *ccoP*) and the flavoprotein oxidoreductases (*gltB*, *gltD*, *gor*, *lpd*, *putA*, *nadB*), while the individually annotated Complex I subunits (*nuoA*–*nuoN*) did not differ. Significance was assessed by Kruskal-Wallis tests corrected by the Benjamini-Hochberg procedure and is indicated on the gene labels: * adjusted $p < 0.05$; ** adjusted $p < 0.01$. Unlabelled genes were not significant.

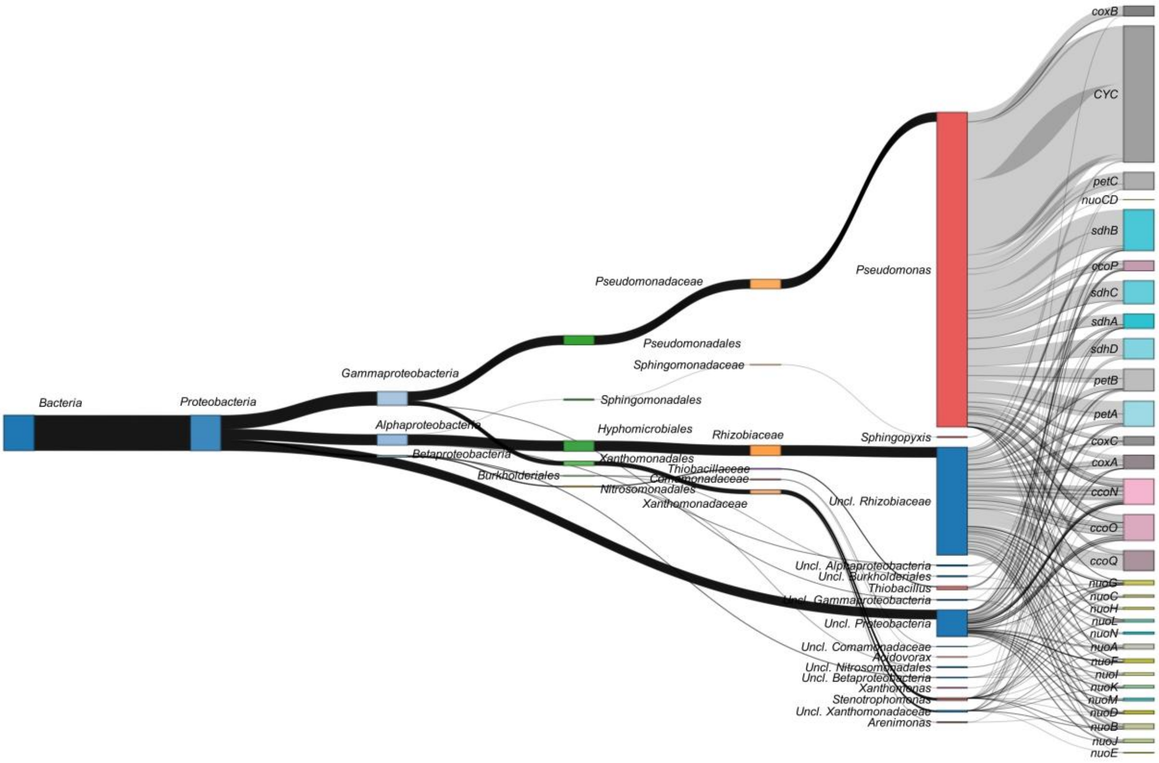


**Figure S4.** Taxonomic origin of respiratory and electron-transport gene expression under high-frequency redox cycling. Sankey diagram of transcript flow for 30 electron-transport and respiratory genes across taxonomic ranks (superkingdom to genus), with hosts assigned from contig taxonomy. Edge width between taxonomic ranks is the number of gene occurrences; genus-to-gene edge width is proportional to mean per-gene TPM.

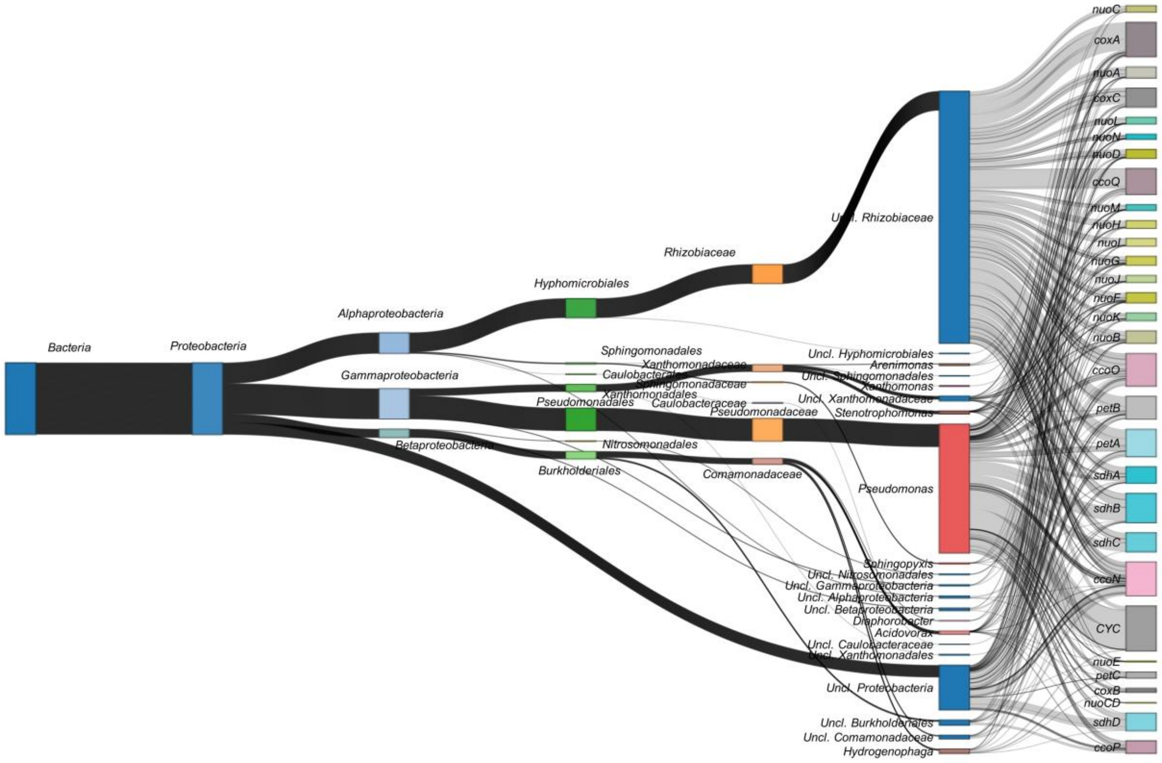


**Figure S5.** Taxonomic origin of respiratory and electron-transport gene expression under medium-frequency redox cycling. Sankey diagram of transcript flow for 30 electron-transport and respiratory genes across taxonomic ranks (superkingdom to genus), with hosts assigned from contig taxonomy. Edge width between taxonomic ranks is the number of gene occurrences; genus-to-gene edge width is proportional to mean per-gene TPM.

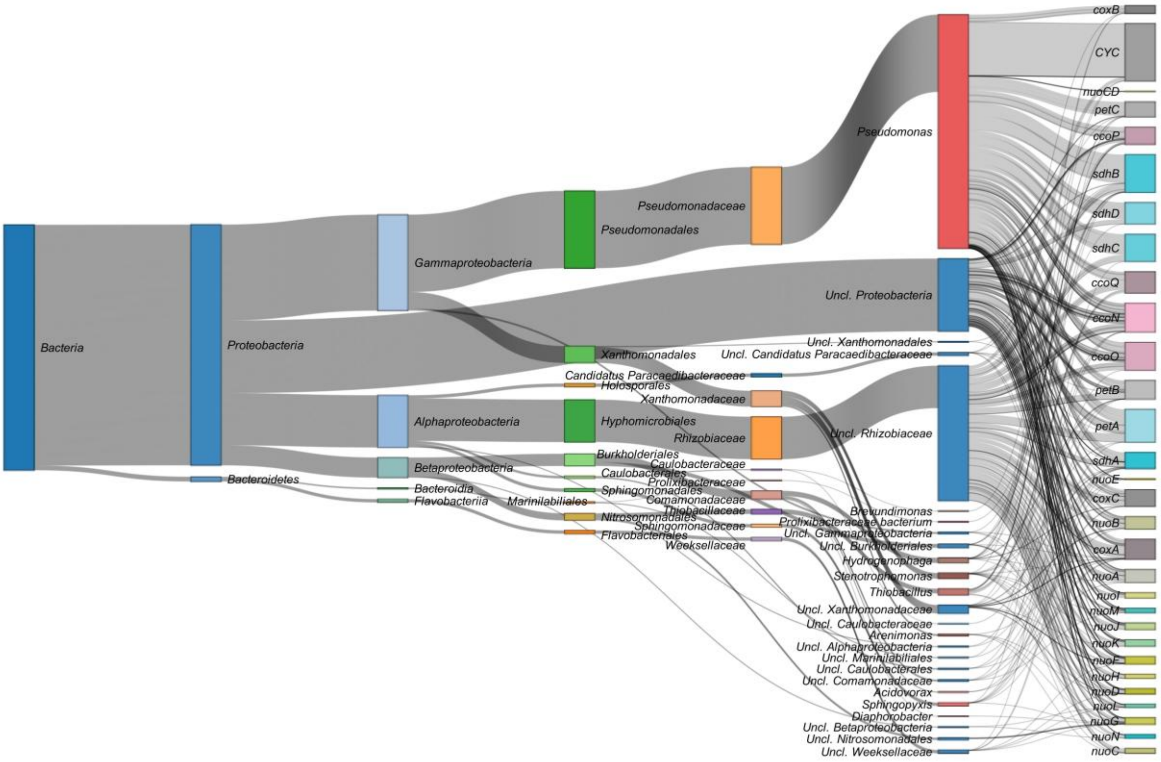


**Figure S6.** Taxonomic origin of respiratory and electron-transport gene expression under low-frequency redox cycling. Sankey diagram of transcript flow for 30 electron-transport and respiratory genes across taxonomic ranks (superkingdom to genus), with hosts assigned from contig taxonomy. Edge width between taxonomic ranks is the number of gene occurrences; genus-to-gene edge width is proportional to mean per-gene TPM.

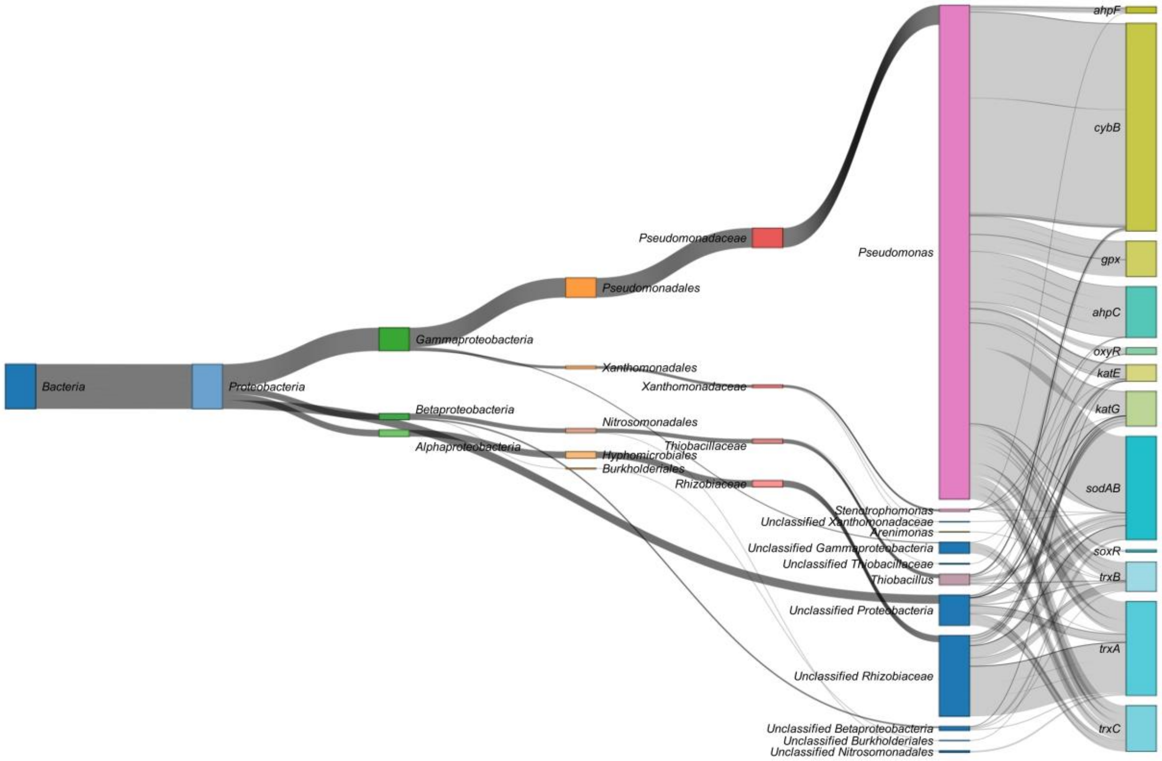


**Figure S7.** Taxonomic origin of oxidative-stress response gene expression under high-frequency redox cycling. Sankey diagram of transcript flow for 12 oxidative-stress and antioxidant defense genes across taxonomic ranks (superkingdom to genus), with hosts assigned from contig taxonomy. Edge width between taxonomic ranks is the number of gene occurrences; genus-to-gene edge width is proportional to mean per-gene TPM.

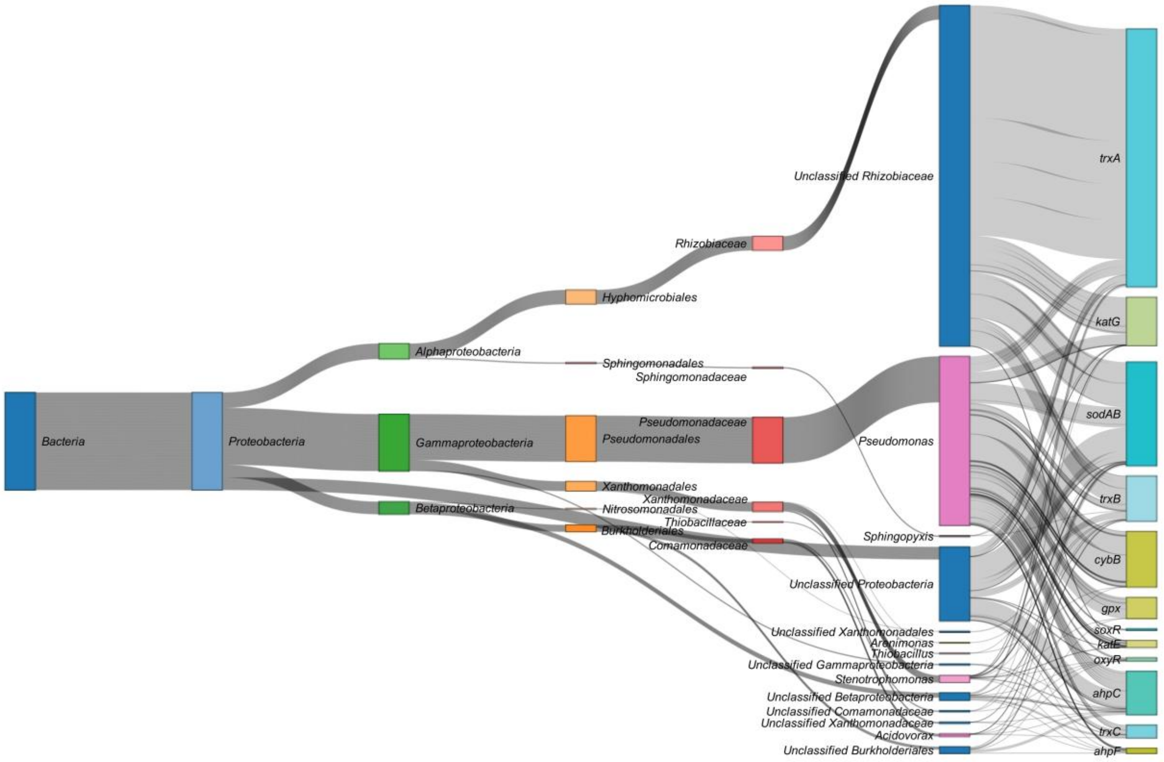


**Figure S8.** Taxonomic origin of oxidative-stress response gene expression under medium-frequency redox cycling. Sankey diagram of transcript flow for 12 oxidative-stress and antioxidant defense genes across taxonomic ranks (superkingdom to genus), with hosts assigned from contig taxonomy. Edge width between taxonomic ranks is the number of gene occurrences; genus-to-gene edge width is proportional to mean per-gene TPM.

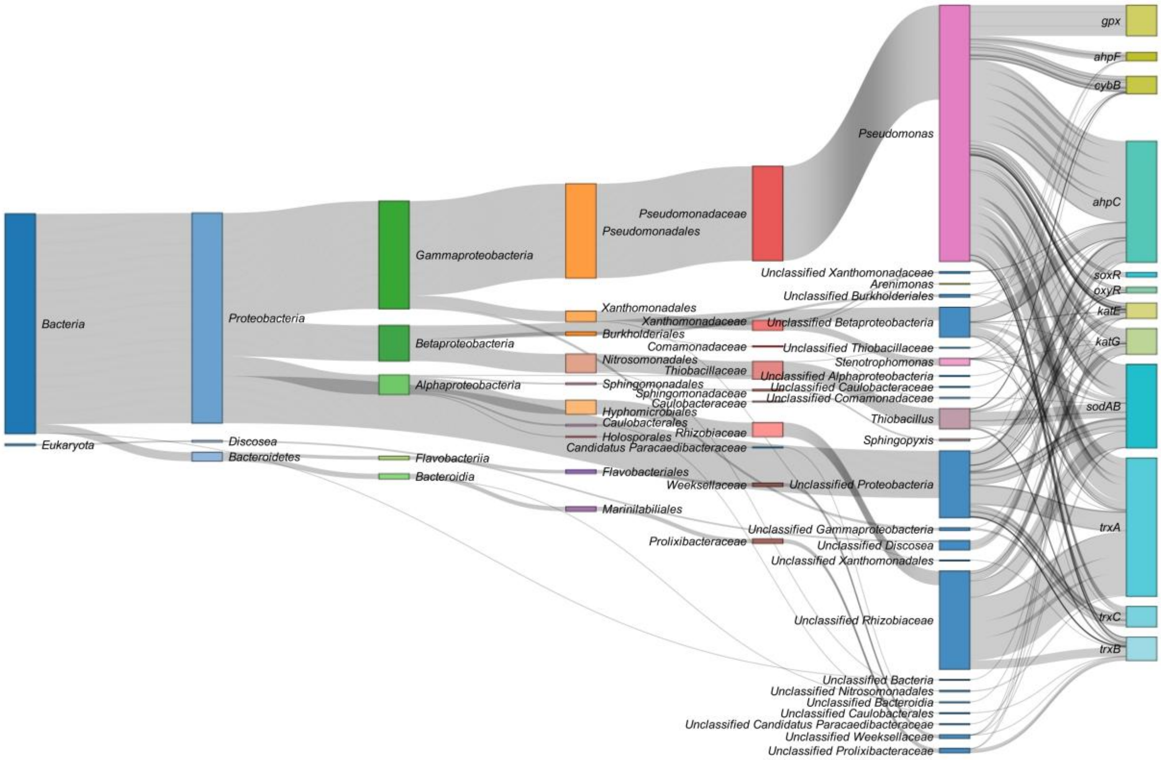


**Figure S9.** Taxonomic origin of oxidative-stress response gene expression under low-frequency redox cycling. Sankey diagram of transcript flow for 12 oxidative-stress and antioxidant defense genes across taxonomic ranks (superkingdom to genus), with hosts assigned from contig taxonomy. Edge width between taxonomic ranks is the number of gene occurrences; genus-to-gene edge width is proportional to mean per-gene TPM.

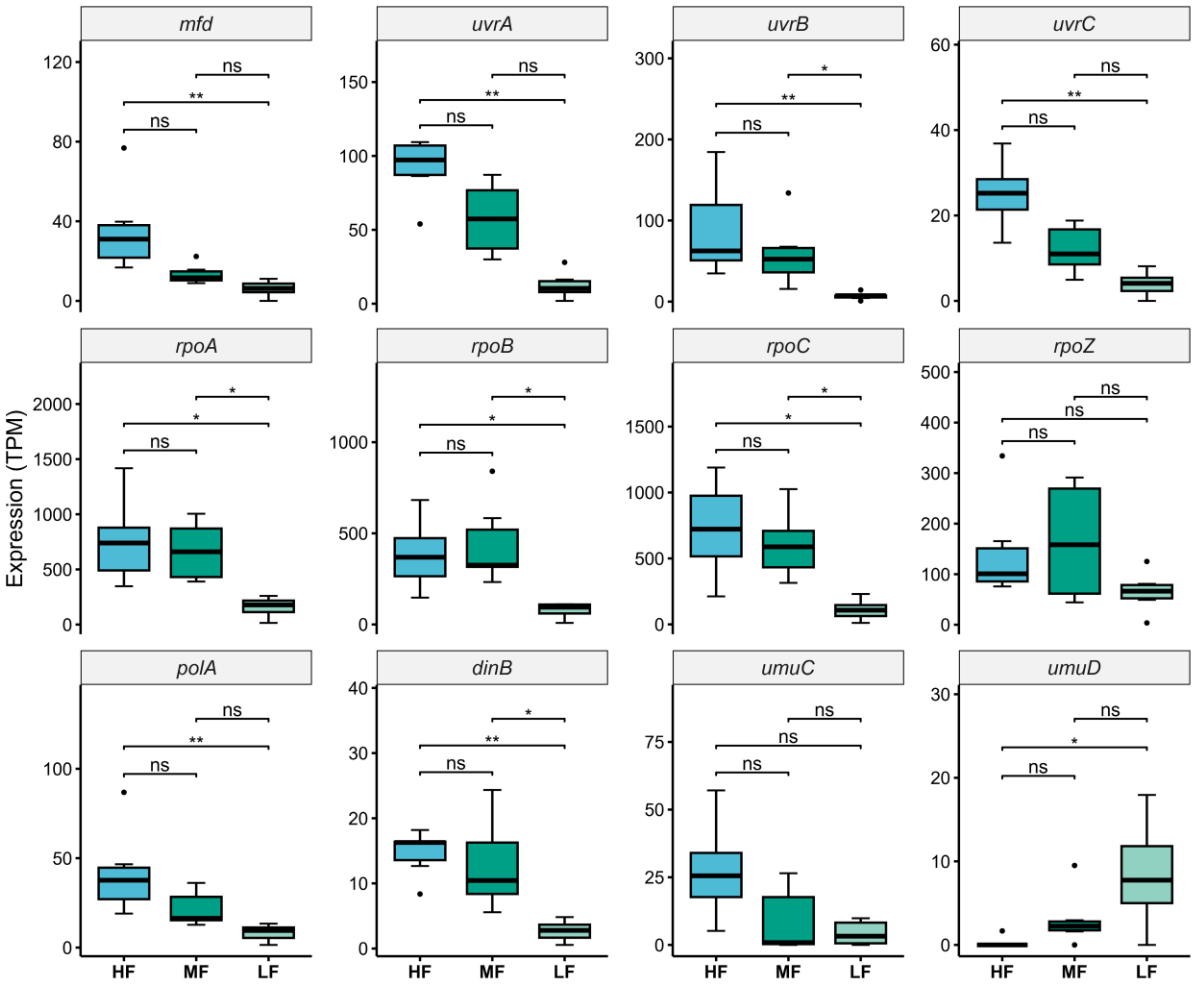


**Figure S10.** Expression of transcription-coupled DNA repair genes across redox fluctuation frequencies. Boxplots of transcript abundance (TPM) for 12 genes of the transcription-coupled repair pathway and the associated RNA polymerase core, under high-frequency (HF), medium-frequency (MF), and low-frequency (LF) oxic–anoxic cycling (n = 6 per treatment). Panels are grouped by function: transcription–repair coupling factor (*mfd*), nucleotide excision repair (*uvrA*, *uvrB*, *uvrC*), RNA polymerase subunits (*rpoA*, *rpoB*, *rpoC*, *rpoZ*), DNA polymerase I (*polA*), and translesion polymerases (*dinB*, *umuC*, *umuD*). Significance was assessed by Kruskal-Wallis tests with Dunn post-hoc contrasts, both corrected by the Benjamini-Hochberg procedure: ns, not significant; * adjusted $p < 0.05$; ** adjusted $p < 0.01$.

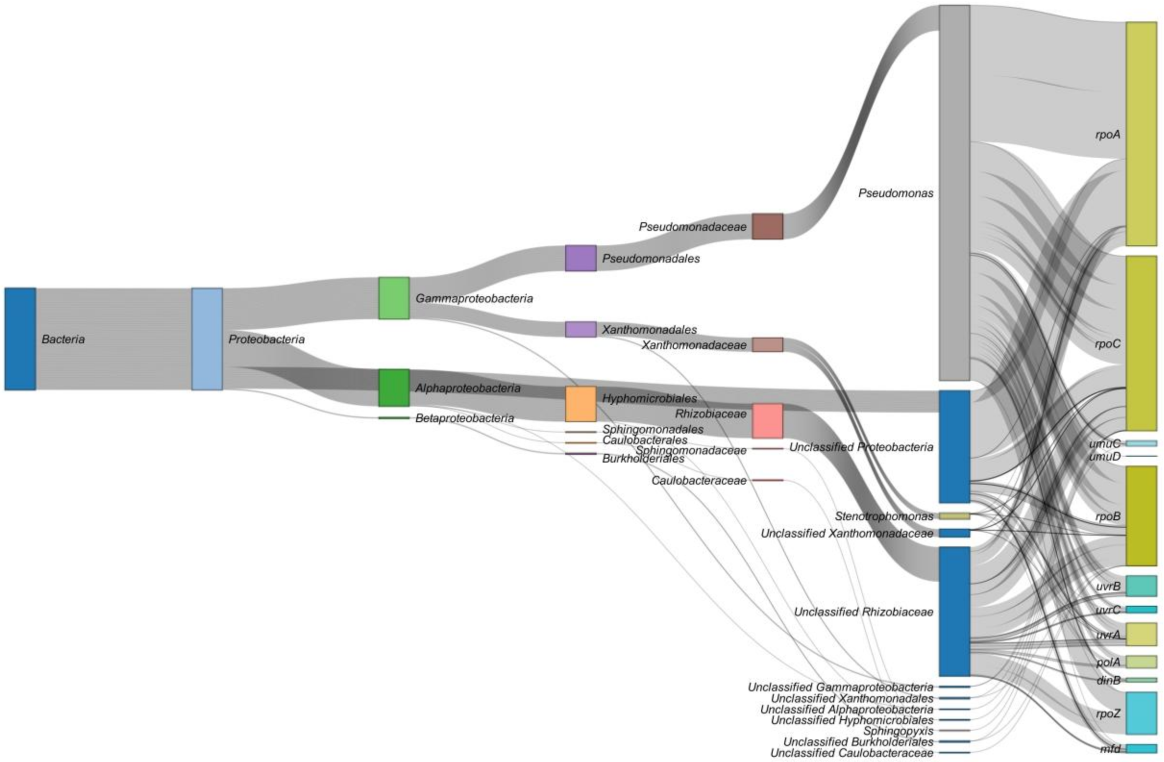


**Figure S11.** Taxonomic origin of transcription-coupled DNA repair gene expression under high-frequency redox cycling. Sankey diagram of transcript flow for 12 nucleotide excision repair and translesion synthesis genes across taxonomic ranks (superkingdom to genus), with hosts assigned from contig taxonomy. Edge width between taxonomic ranks is the number of gene occurrences; genus-to-gene edge width is proportional to mean per-gene TPM.

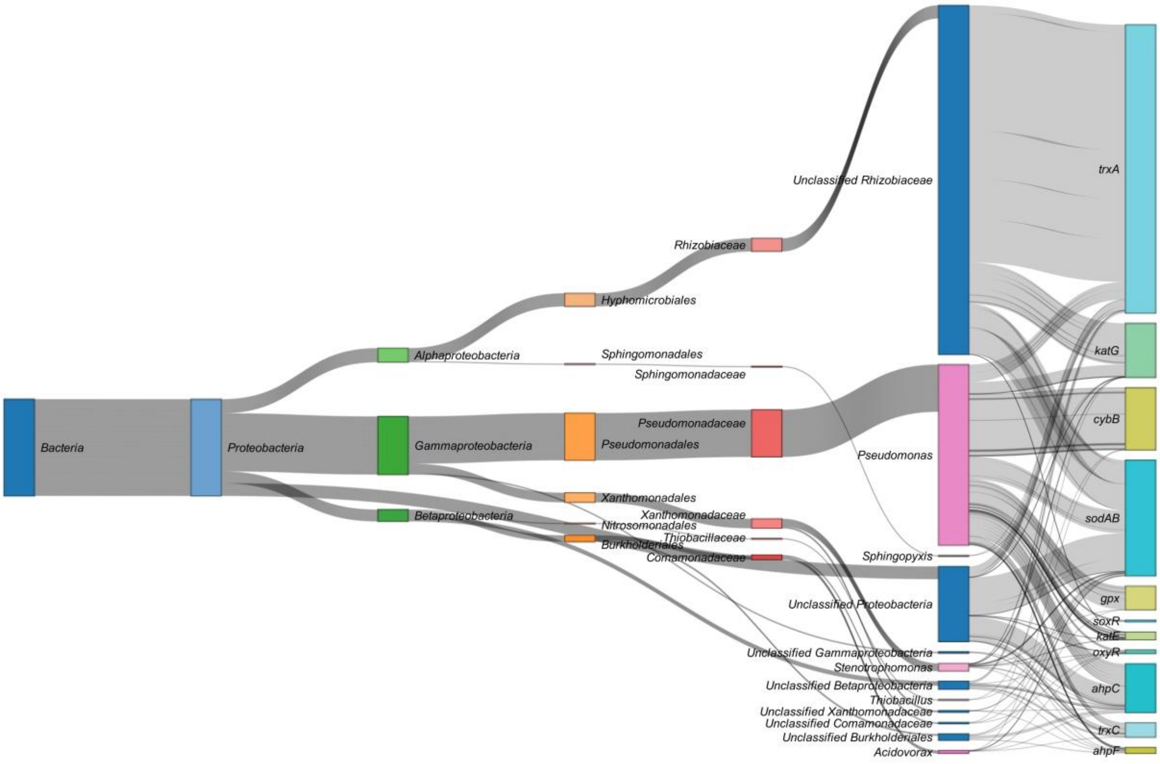


**Figure S12.** Taxonomic origin of transcription-coupled DNA repair gene expression under medium-frequency redox cycling. Sankey diagram of transcript flow for 12 nucleotide excision repair and translesion synthesis genes across taxonomic ranks (superkingdom to genus), with hosts assigned from contig taxonomy. Edge width between taxonomic ranks is the number of gene occurrences; genus-to-gene edge width is proportional to mean per-gene TPM.

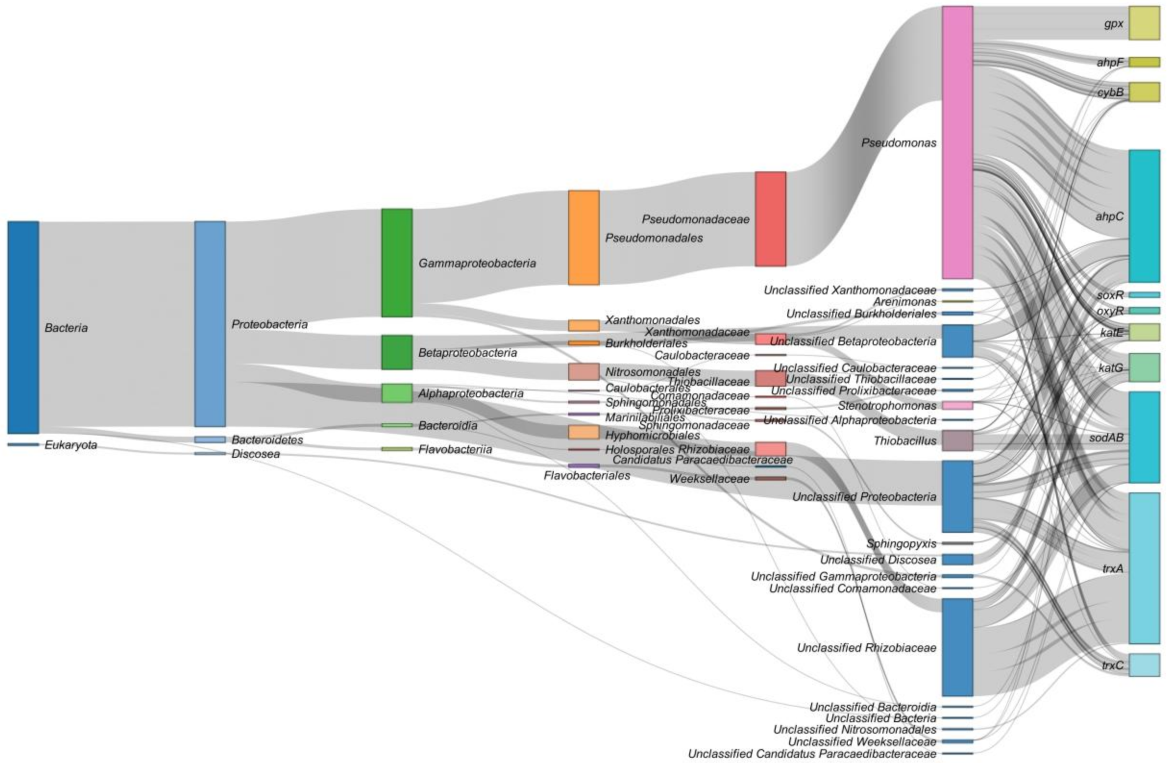


**Figure S13.** Taxonomic origin of transcription-coupled DNA repair gene expression under low-frequency redox cycling. Sankey diagram of transcript flow for 12 nucleotide excision repair and translesion synthesis genes across taxonomic ranks (superkingdom to genus), with hosts assigned from contig taxonomy. Edge width between taxonomic ranks is the number of gene occurrences; genus-to-gene edge width is proportional to mean per-gene TPM.

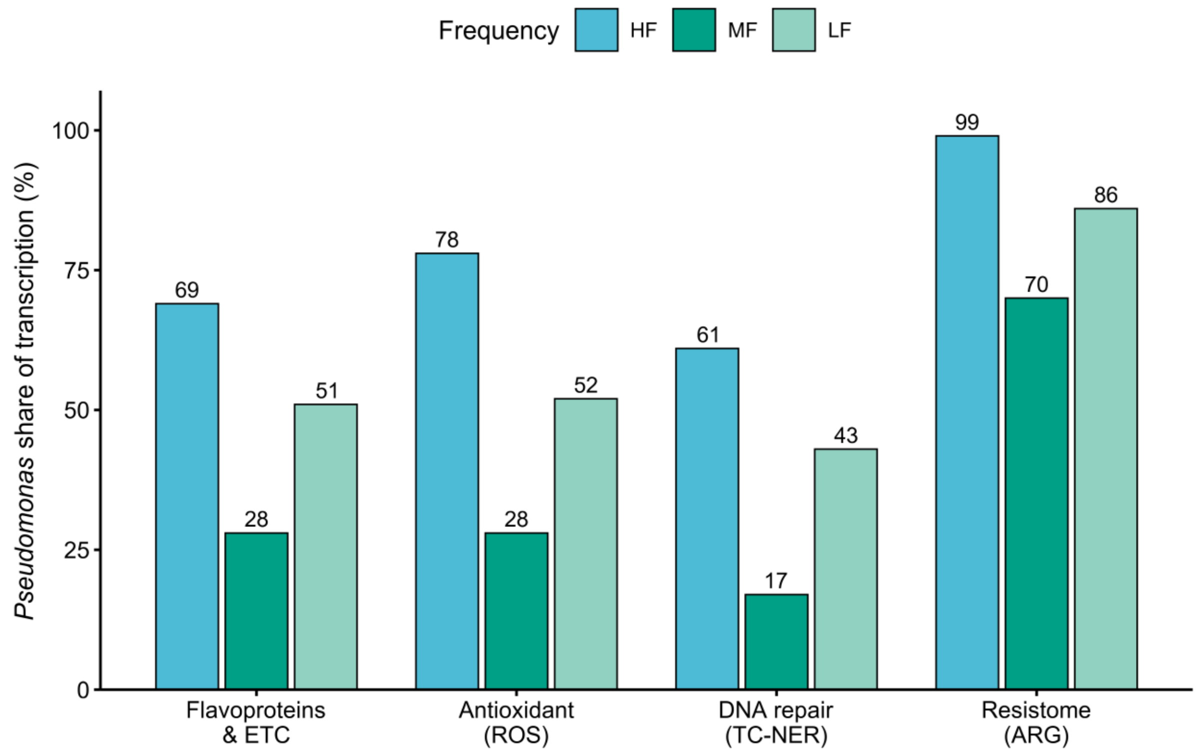


**Figure S14.** *Pseudomonas* share of transcription across functional layers and frequencies. Fraction of transcription assigned to *Pseudomonas* for four gene sets (flavoprotein/ETC, antioxidant, DNA repair, and resistome) under HF, MF, and LF cycling.

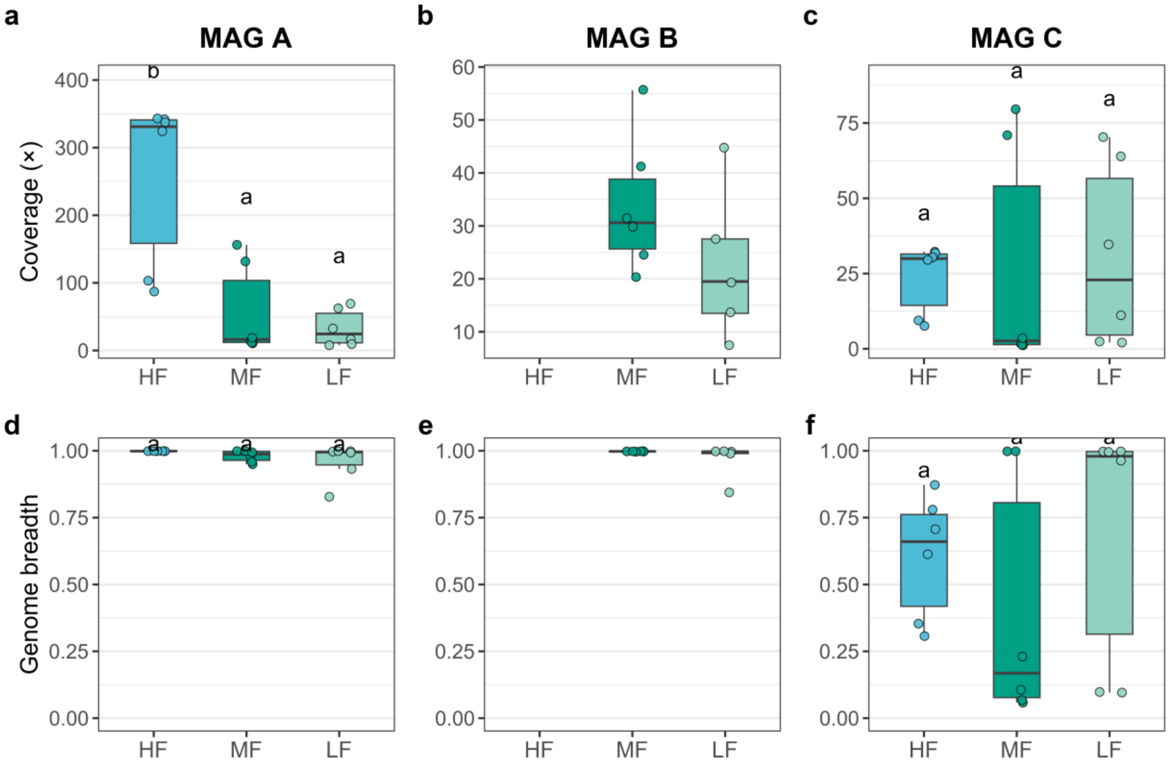


**Figure S15.** Recovery of the three Pseudomonas MAGs across redox cycling frequencies. Coverage (a–c) and genome breadth (d–f) for MAG A, B, and C under high- (HF), medium- (MF), and low-frequency (LF) cycling (n = 6, except MAG B under LF, n = 5). MAG B was absent from all HF replicates; MAG C was recovered at low breadth under HF (median 66%) and MF (17%). Only MAG A reached near-complete breadth (>98%) in every treatment, justifying its use for the strain-resolved analysis. Boxes show the interquartile range and median; shared letters indicate no significant difference (Kruskal–Wallis with Dunn, BH-corrected).

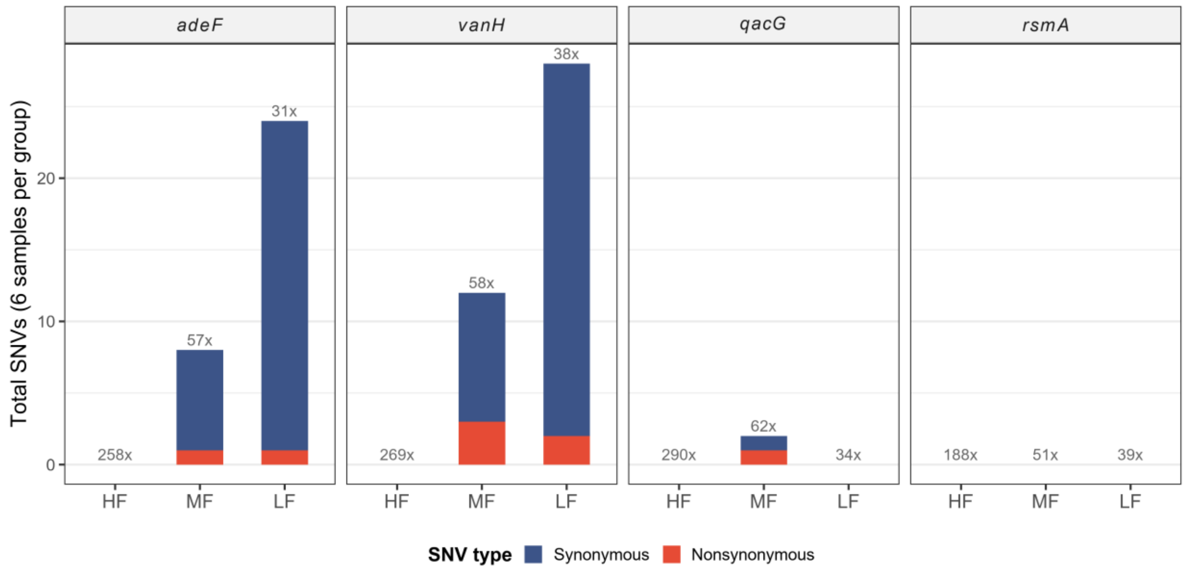


**Figure S16.** Resistance genes in Pseudomonas MAG A vary only under MF and LF cycling, almost entirely at synonymous sites. Total SNVs summed across the six replicates per treatment for the four MAG A ARGs (*adeF*, *vanH*, *qacG*, *rsmA*) under high- (HF), medium- (MF), and low-frequency (LF) cycling, split into synonymous (blue) and nonsynonymous (red). No ARG carried an SNV under HF despite the deepest coverage; MF and LF accumulated 30 and 76 SNVs (24 of 30 and 72 of 76 synonymous). Numbers above bars are mean per-sample coverage (×).
